# Visualizing scRNA-Seq Data at Population Scale with GloScope

**DOI:** 10.1101/2023.05.29.542786

**Authors:** Hao Wang, William Torous, Boying Gong, Elizabeth Purdom

## Abstract

Increasingly, scRNA-Seq studies explore cell populations across different samples and the effect of sample heterogeneity on organism’s phenotype. However, relatively few bioinformatic methods have been developed which adequately address the variation between samples for such population-level analyses. We propose a framework for representing the entire single-cell profile of a sample, which we call a GloScope representation. We implement GloScope on scRNA-Seq datasets from study designs ranging from 12 to over 300 samples and demonstrate how GloScope allows researchers to perform essential bioinformatic tasks at the sample-level, in particular visualization and quality control assessment.

## 1 Background

Single-cell sequencing data has the potential to considerably enhance our comprehension of human health demonstrating how individual cell differences affect disease outcomes. Initially, single-cell sequencing studies examined the scope of cell diversity found in biological systems, including large projects such as the Human Cell Atlas Project. Such studies generally obtain large numbers of cells from few individual donors and focus on the shared cell type diversity. However, an increasing number of scRNA-Seq investigations target patient populations and emphasize the impact of single-cell variation on human health outcomes. These population-based scRNA-Seq studies typically involve scRNA-Seq data from larger cohorts of individuals who are selected from populations exhibiting various health-related phenotypes.

Despite the plethora of methodological advancements in scRNA-Seq, most current tools were designed for the goal of understanding the single cell level information and lack appropriate strategies for analyzing scRNA-Seq population studies. Most of the current analyses of population scRNA-Seq data tends to consider the individual cells as the primary data unit. Existing tools that do account for population variability focus on identifying individual genes with differential expression [1–3]. Beyond differential expression analysis, sample-level analysis that exist are generally limited to comparisons of the relative proportions of different cell-types between groups of samples [4]. We propose an analysis paradigm that uses the entire single-cell profile of a sample instead of focusing on cells as units. We refer to such an approach as a sample-level (or patient-level) analysis.

Our proposal is based on representing each sample as a distribution of cells. More specifically, we summarize each sample with a probability distribution describing the distribution of cells and their gene expression within the sample. Such a representation allows us to summarize the entire scRNA-profile of a sample into a single mathematical object. In this way, we synthesize the entire single-cell profile of an individual sample while maintaining information regarding the variability of the single-cells. This global representation, which we call GloScope, can be used in a wide variety of downstream tasks, such as exploratory analysis of data at the sample-level or prediction of sample phenotypes. Moreover, this representation does not require classification of sequenced cells into specific cell-types (e.g. via clustering), and therefore is not sensitive to any auxiliary cell-type identification procedure.

We apply the GloScope representation on a variety of published data collected on sample cohorts and demonstrate how the GloScope representation allows for visualization of important biological phenotypes and aids in detection of sample-level batch effects.

## 2 Results

### 2.1 Overview of the GloScope Representation

If we consider trying to model individual samples, we see that the format of scRNA-Seq data when considered as data on samples (not cells) is non-standard. Most computational strategies assume each sample is measured on a shared set of features. Instead, for each sample *i* we observe a matrix *X_i_ ∈ R^g×mi^*, containing the gene expression measurements of that sample across all cells (*g* corresponds to the number of genes and *m_i_* to the number of cells sequenced from sample *i*). There is no direct correspondence between the *m_i_* cells in sample *i* with the *m_j_* cells of sample *j* so there is no immediate way to align data from different samples as input into a statistical model or predictive algorithm.

We propose to create a representation of each sample that does not require explicitly aligning individual cells across samples, but leverages the nature of the observed data to represent each sample in a similar space. We consider the gene measurements for each of the *m_i_* cells to be a sample from the full population of all cells of each sample. The full population of cells defines a probability distribution we designate as *F_i_* on *R^g^*. *F_i_* is a representation of the sample’s entire single-cell profile across all cells and importantly is a mathematical object that can be compared across samples. We do not observe *F_i_*, but we do observe *m_i_* samples from this distribution (the sequenced cells), allowing us to estimate *F_i_* from the data. Thus, we transform each sample from the matrix *X_i_* of observed gene expression measurements to an estimate of the sample’s distribution, *F*^^^*_i_*.

However, because gene expression data lie in a high dimensional space, with the number of genes *g* in the thousands, estimating *F_i_* directly from the cells is intractable. Thus, we assume that there exists a lower dimensional representation or latent variable in *R^d^* which governs the gene expression of a sample. We instead estimate the distribution of this latent variable. We do this by first estimating a lower dimensional representation of our all our cells, for example via methods like PCA or scVI [5] applied to all the cells. This results in a matrix of reduced representation *Z_i_ ∈ R^mi×d^* corresponding to the new coordinates of each cell in this reduced space. We then estimate the distribution *F*^^^*_i_* from the *m_i_* cells in this reduced space.

Unlike the *X_i_*, which have different, unrelated, dimensions for each sample *i*, the *F*^^^*_i_* lie in the space of distributions on *R^d^* and can be compared. As probability measures, these representations are now familiar mathematical objects and sample-level analysis can be done in the space of probability measures. There are many well-known metrics defined on the space of probability measures, such as the Wasserstein distance, and downstream analysis can be performed after choosing a metric to quantify pairwise sample differences. We call this representation of samples the GloScope representation, and we illustrate this transformation in Fig. 1. For our examples, we use the square root of the symmetrized Kullback-Leibler (KL) divergence to quantify the differences between sample distributions; while not a proper metric, this divergence can be effectively used to create a global representation of probability distributions [6] (see Methods for details).

**Fig. 1.**
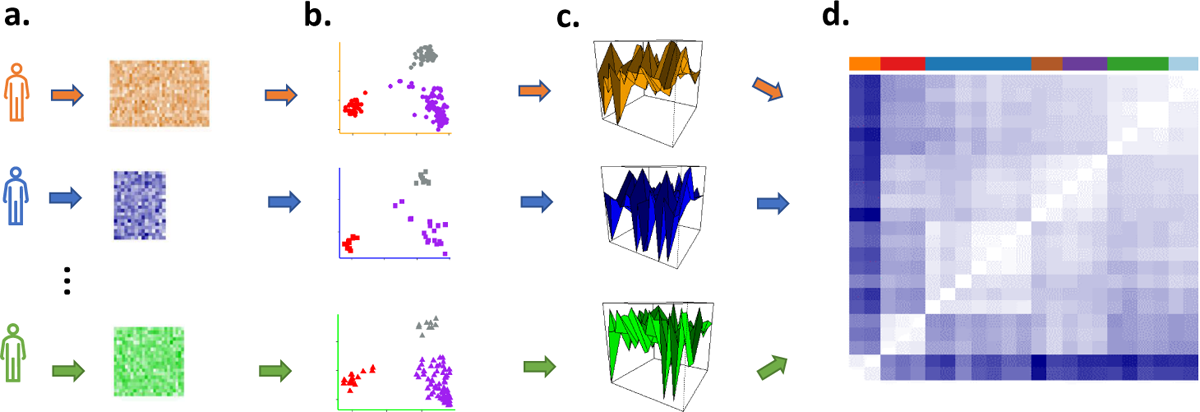
Illustration of the GloScope representation of a sample’s scRNA-Seq data matrix. *X_i_* **as a distribution** *F*^^^*i*. (a) Each sample contributes a *g × m_i_* matrix of gene expression values. (b) A lower dimensional latent representation is estimated across all cells and samples, resulting in each cell being represented in a lower-dimensional space (c) GloScope estimates the distribution *F*^^^*i* for each sample, and then (d) calculates the statistical divergence between each pair of samples, *d*(*F*^^^*i*, *F*^^^*j*).

The resulting pairwise-divergences can be used by many standard statistical or machine learning methods. We primarily concentrate in this work on the use of the GloScope representation for the purpose of visualization and exploratory data analysis. The pairwise divergences between GloScope-represented samples can be given as input to canonical divergence analysis methods such as Multidimensional Scaling [7], which creates coordinate system to represent the samples that capture the pairwise divergences. We will demonstrate that such a visualization enables detection of possible batch effects or outliers and exploratory assessment of the strength of phenotypic differences between our samples. We can also use the divergences to numerically quantify the separation of groups of samples using silhouette width or ANOSIM statistics (see Methods). This allows us to quantify how separated samples are due to a biological condition of interest (e.g. healthy vs diseased samples), or alternatively how separated samples are due to a design artifact (e.g. different processing centers). Beyond EDA, our representation can also be used for other important downstream tasks, include clustering of samples, global hypothesis tests for differences between sample populations, and prediction of phenotypes (for example via kernel prediction methods, e.g. Hofmann et al. [8], Wang et al. [9]).

### GloScope in the scRNA-Seq Pipeline

There are many existing methods for working with scRNA-Seq data, and GloScope is designed to fit into standard pipelines and complement existing quality-control and EDA strategies. GloScope takes as input low-dimensional latent representations of the individual cells, which can come from standard embeddings of the original cell data, like PCA, scVI; from batch-correction methods like Harmony [10]; or from integration methods that harmonize data processed on different gene definitions [see 11, for a review]. GloScope can be performed at different stages of the pre-processing, allowing checks at each stage of whether patient-level artifacts, like processing batches, are inappropriately contributing to differences in the samples.

### Using Cell-type Composition

Our GloScope approach to creating a global representation uses the entire gene distribution *F_i_*, which encodes both cell-type composition and gene expression. However, the underlying logic of GloScope could also be applied to compare only cell-type composition. Specifically, if each cell can be classified into one of *K* subtypes, then we observe for each sample the proportion of cells in each cell-type, *π*^*_i_* = (*π*^*_i_*_1_*, …, π*^*_iK_*) *∈ R^K^*. *π*^*_i_* is an estimate of a probability distribution, only now a simpler discrete distribution into *K* groups. We can use the GloScope strategy in a similar way to globally compare samples, only now restricted to only differences in cell-type composition. Comparison of cell-type composition has been proposed for globally comparing single-cell samples [12–16], and there has been some limited work in analysis of data from flow-cytometry using cell-type compositions to globally compare samples which has similarities to using GloScope on the proportions [12, 17–19]. Unlike a full GloScope representation, applying GloScope on the cluster proportion vector requires classifying cells into subtypes before application of the method. Accurate identification of cells into subtypes is often a manual and time-consuming process, which makes this approach less useful for the exploratory data analysis that is often upstream of the subtype identification step (see Section 2.5). However, GloScope applied to the clusters can be used for more formal hypothesis testing of significant global differences in cell-type composition. In what follows, we will refer to GloScope applied to the vector cluster proportions as GloProp, as opposed to our standard implementation which calculates an estimate of the full gene expression density.

### 2.2 Visualization of patient and sample phenotypes using GloScope representations

In this section we demonstrate the utility of the GloScope representation to visualize and evaluate sample-level phenotypic differences. As an initial illustration, we consider two datasets with replicate samples collected for each phenotype, where the phenotypes have well-known biological differences in cell-type structure. These serve as an initial proof-of-concept of the GloScope representation.

The first dataset is scRNA-Seq data from the mouse cortex [20]. Here the samples are cells from different regions of the brain with replication in each from three genetically identical mice. This is a dataset where we know the regions have distinct compositions of cell types and gene expressions. When we visualize these samples using the GloScope representation in Fig. 2A, we see these distinctions clearly. The samples from the two main subdivisions of the cortex, isocortex (CTX) and hippocampal formation (HPF), clearly separate. Furthermore, we see that replicate samples from the same region strongly cluster with each other, while different regions are generally well separated. Within the CTX region, we observe blocks of biologically meaningful brain region groups such as the sensory and visual area: primary somatosensory (SSp), posterior parietal association (PTLp), visual area (VIS), and the Somatomotor areas: primary motor (MOp) and secondary motor (MOs). We also observe clustering of physically adjacent brain regions such as temporal association, perirhinal, and ectorhinal areas (TEa-PERI-ECT), agranular insular (AI), prelimbic, infralimbic, orbital area (PL-ILA-ORB) and anterior cingulate (ACA).

**Fig. 2.**
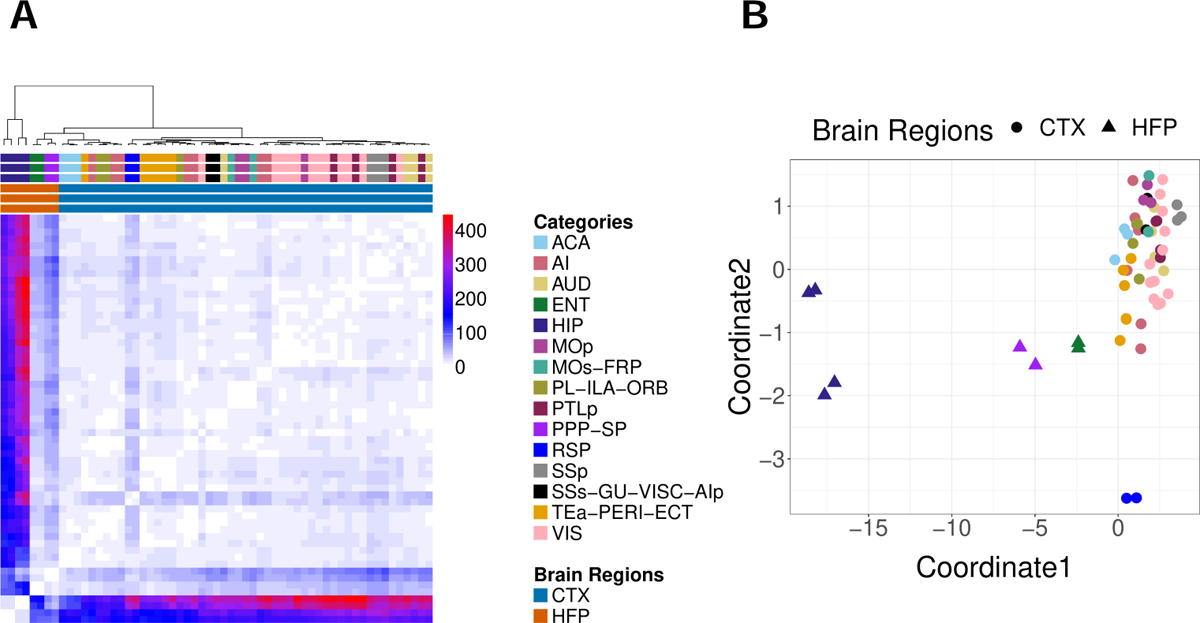
Demonstration of the GloScope representation on 59 mice samples. [20]. (A) Heatmap representation of the estimate of the divergences between the samples based on the GloScope representation. (B) A two dimensional representation via MDS of the divergences shown in (A) GloScope used the GMM estimate of the density in the first 10 PCA dimensions. The individual regions represent subregions of two main divisions of the cortex: the isocortex (CTX) and hippocampal formation (HPF). HPF is further divided into hippocampal region (HIP), and the retrohippocampal region (RHP) which is represented by the entorhinal region (ENT) and the remaining RHP, a joint dissection region of postsubiculum (POST)-presubiculum (PRE)-parasubiculum (PAR) region, subiculum (SUB), and prosubiculum (ProS) region (i.e, PPP-SP). The remaining regions are divisions of the CTX.

Next we consider skin cell samples from a study of twelve patients, Cheng et al. [21] consisting of nine healthy skin samples from the foreskin, scalp, and trunk alongside three inflamed skin samples collected from truncal psoriatic skin. We expect marked differences between cellular distributions collected at the different locations in the body due to varying proportions of cell types in certain tissues. For instance the authors note different types of main basal keratinocytes and melanocytes dominate in scalp and trunk samples, as compared to foreskin tissues. Our visualization of the GloScope representations of this data in Fig. 3 shows a clear clustering of skin samples collected from similar locations on the body, and a separation of both the foreskin and psoriasis samples from scalp and trunk samples, echoing the conclusions of the authors who identified a keratinocyte subpopulation which separates these phenotypes from the scalp and trunk control samples Cheng et al. [21].

**Fig. 3.**
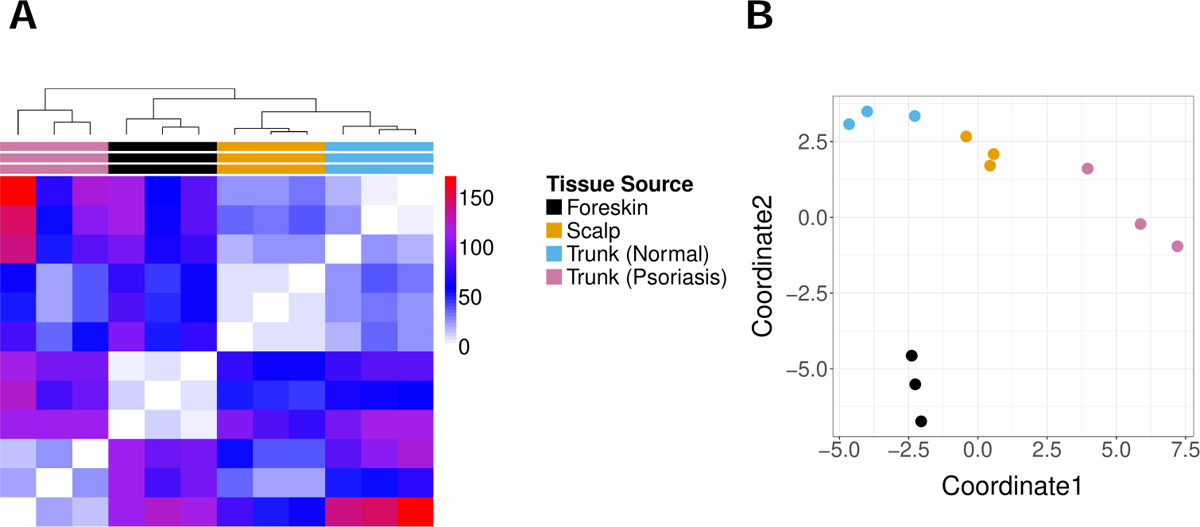
GloScope representation of 12 skin rash patients collected in various locations and conditions in. [21]. (A) A heatmap visualization of the estimate of the symmetrized KL divergence between the samples’ GloScope representation. (B) A two dimensional MDS representation of the divergences. The divergences were calculated using the GMM density estimation based on PCA estimation of the latent space in 10 dimensions.

Next we demonstrate the GloScope representation on additional datasets of patient cohorts where the samples are patients with differing disease phenotypes: 1) COVID lung atlas data from [22], which contains 27 samples, either diagnosed with COVID-19 or healthy control samples, and 2) Colorectal cancer data with 99 samples (after quality control), grouped into three phenotypes: healthy, mismatch repair-proficient (MMRp) tumors, and mismatch repair-deficient (MMRd) tumors [23]. The use of GloScope on these datasets demonstrates its utility for the visualization of both sample and phenotype variability. For the COVID lung samples (Fig. 4A), we can easily see the separation between COVID-infected and healthy donors, matching the observation of Melms et al. [22] that lung samples from COVID patients were highly inflamed. For the colorectal cancer data, visualization of the GloScope representation shows healthy samples well separated from the tumor samples (Fig. 4B). Though the two types of tumors do not separate in this visualization, an Analysis of Similarities (ANOSIM) test of significance [24] applied to their GloScope divergences between these two groups does find their representations to be significantly different (*p* = 0.001), indicating that the representation is encapsulating systematic differences between the two tumors (see Methods section).

**Fig. 4.**
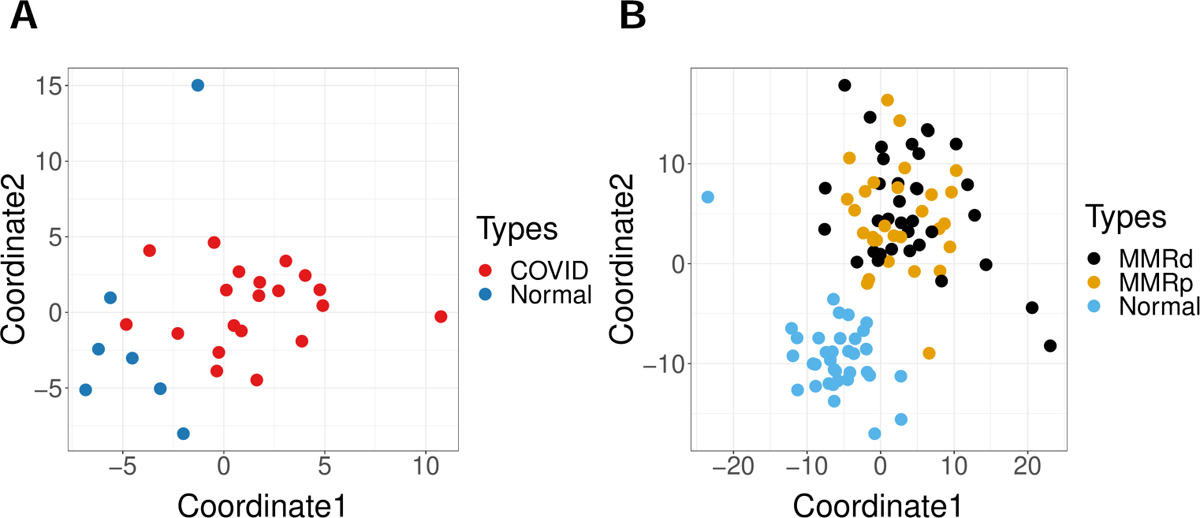
Examples of MDS plot of the dissimilarities calculated from GloScope representation. (A) 27 samples of COVID lung atlas data that are either healthy samples of COVID patients from Melms et al. [22]; (B) 99 colon samples from mismatch repair-proficient (MMRp) tumors, mismatch repair-deficient (MMRd) tumors and healthy samples from Pelka et al. [23]. The dissimilarity matrices were calculated using the GMM density estimate based on PCA estimates of the latent space in 10 dimensions.

### 2.3 Quantitative Evaluation of GloScope via Simulation

We use simulation experiments to quantify GloScope’s efficacy at detecting various classes of single-cell differences that might be observed due to differences in samples’ phenotype. We simulate sample-level data where different aspects of the single-cell composition of a sample vary depending on their group assignment; for simplicity we consider only two different phenotypic groups. Count matrices were generated from a pipeline modified from that presented in the R package muscat [1] (see Methods for details).

We focus on two basic biological scenarios that could causes phenotypic-based dissimilarity between scRNA-Seq samples which we would want the GloScope representation to accurately reflect: differential cell-type composition and differential gene expression. By cell-type composition, we refer to the proportion of various cell-types found in a sample; for example an inflammatory disease phenotype might result in a higher proportion of immune cells in the patient than in a healthy sample. Cell-type gene expression differences (DE) refers to differences across samples in the marginal gene expression levels within cells of a certain type. For example the IL2 gene has more expression within the T-cells of inflammation tissue samples when compared to the its expression in T-cells of healthy samples. Both types of differences are biologically plausible and can co-exist. We also note that in practice the distinction between these two can blur: many genes exhibiting sufficiently strong differential expression between phenotypes will result in the creation of a novel cell-type for all practical purposes, thereby corresponding to differential cell-type composition and vice versa.

In our simulations we evaluate how well these two types of differences are detected by GloScope. We create datasets demonstrating either differentially expressed genes or differential cell-type composition. We see that the average differences between samples in different different phenotype groups, as measured by our GloScope representation, appropriately increase in response to both increased differences in global cell composition (Fig. 5A) and increased differential gene expression (Fig. 5B). This indicates that our representation effectively reflects both types of changes. Similarly, when increased sample variability is added, both in global cell composition and gene expression, our GloScope representation correspondingly shows increased within-group variability (Fig. 5C and 5D).

**Fig. 5.**
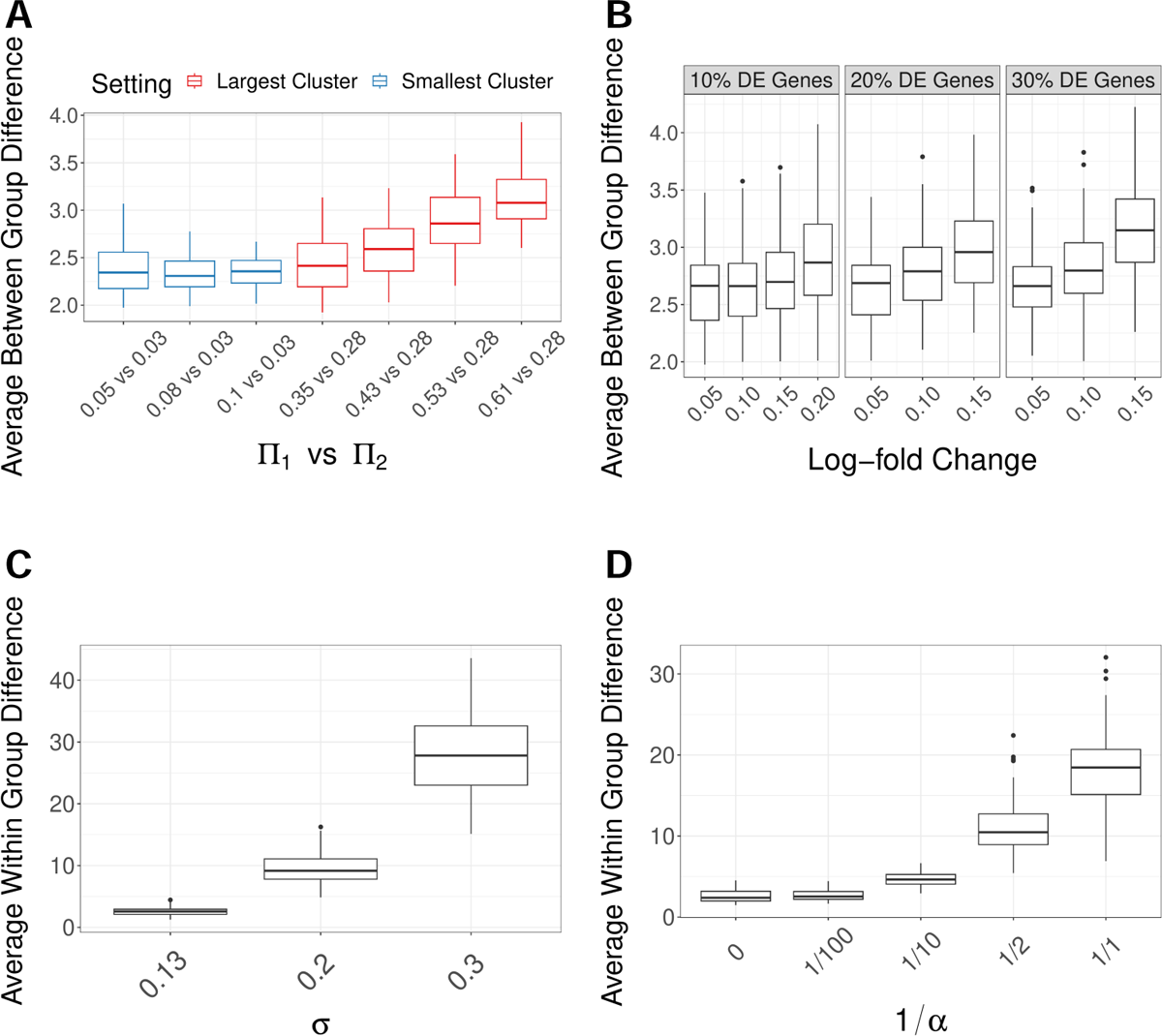
GloScope captures simulated effects. (A) and (B) show how the average GloScope divergence between samples in different phenotype groups increases with (A) increased cell composition differences and (B) increased gene expression differences. The cell composition differences in (A) are color-coded as to whether the major changes were in the two groups’ largest cluster or smallest cluster (the actual values of the proportion changes in the largest or smallest group, Π_1_ vs Π_2_, are labeled in the legends). Plots (C) and (D) shows how the average GloScope divergence between samples in the same phenotype group increases with (C) increased sample variability in gene expression differences and (D) increased cell composition differences. All boxplots show these averages over 100 simulations. The dissimilarity matrices were calculated using the GMM-based GloScope representation based on PCA estimates of the latent space in 10 dimensions. For choices of kNN with scVI or PCA and GMM with scVI, see Additional File 1: Fig S7-S10

We can use our GloScope representation to compare different choices of the design or analysis of the experiment, based on how well the two phenotypic groups separate in the GloScope representation. To do so, we perform analysis of similarities (ANOSIM), a hypothesis test for differences between groups based on observed pairwise divergences on samples [24]. ANOSIM takes as input divergences between samples and tests whether divergences are significantly larger between samples in different groups compared with those found within groups based on permutation testing (see Methods for more details). Evaluation of ANOSIM over many simulations gives the power of the test in different settings, resulting in a metric to compare choices in our analysis.

Using these power computations, we can see that changes in the sample variability and sample size are reflected as expected in these power calculations: increasing all of these sources of variability naturally reduces the power (Additional File 1: Fig. S1). These types of simulations, in conjunction with our GloScope representation, can be used to evaluate design choices at the sample-level, such as the number of samples needed to reach a desired power level. Unsurprisingly, differences in cell-composition in large clusters are more easily detected than similar differences in small clusters (Fig. 6A), and gene expression differences concentrated in small clusters are harder to detect than those found in large clusters (Fig. 6C).

**Fig. 6.**
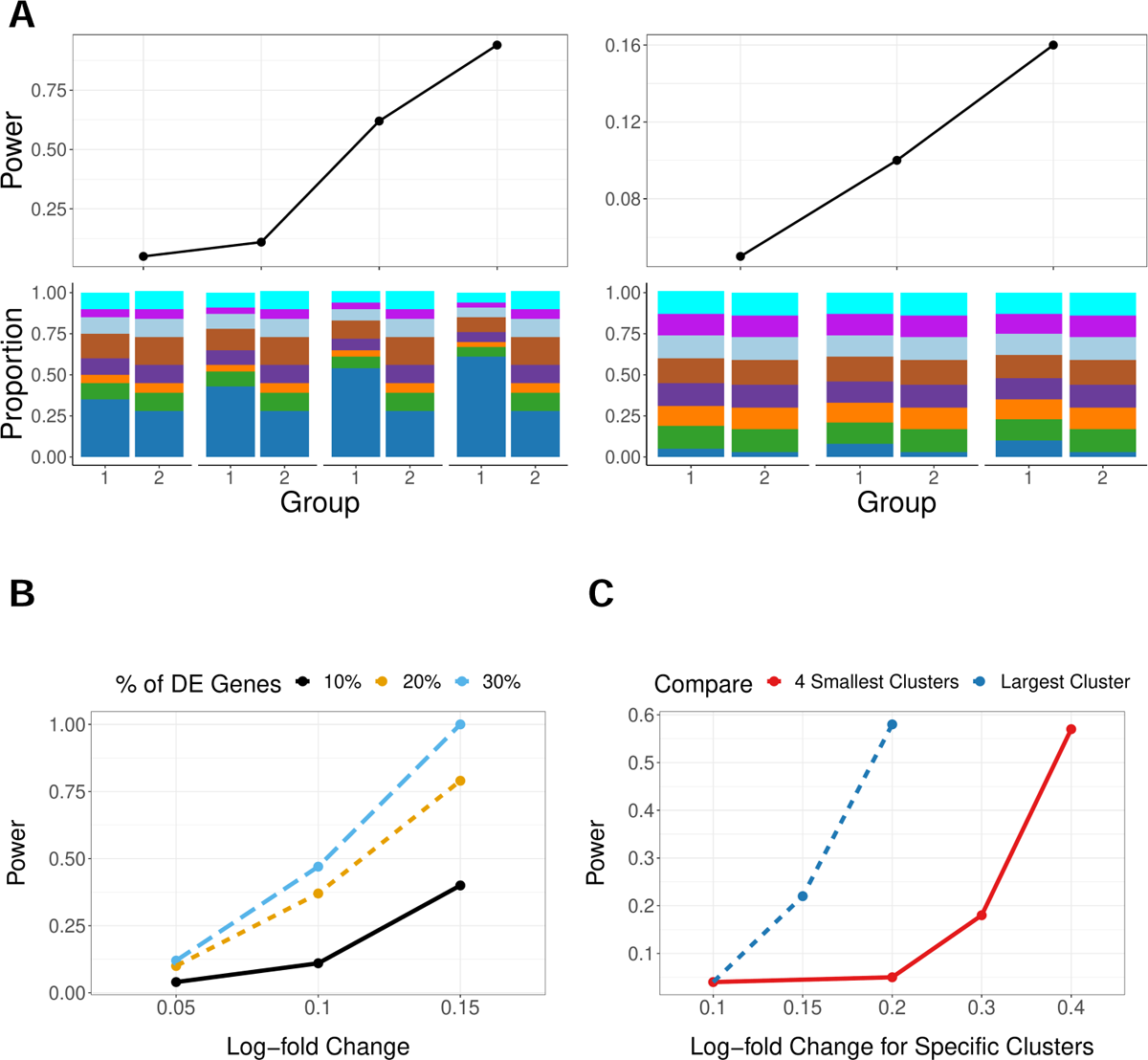
ANOSIM power on simulated data (y-axis) under different conditions. (A) Changes in only the cell-type composition (no DE genes), with major changes in the two groups’ largest cluster (left) or smallest cluster (right). The cell-type composition is visualized in the lower panels. (B) Increasing percentage of DE genes (*ρ_DE_*) with average log-fold change changing from 0.05, 0.1, and 0.15 (x-axis). (C) Changes of log-fold-changes concentrated in specific cell-types/clusters (*ω_k_*), quantified as relative to the baseline log-fold change *θ* =0.05; the two lines correspond to whether the log-fold changes were in the largest cluster (representing *π_k_* = 40% proportion of cells) or for the 4 smallest cluster (representing *π_k_*= 30% proportion of cells). Power calculations were done on relatively small groups to show the full range of changes (n=10 samples in each group) with *m* = 5, 000 cells per sample; the sample level variability parameter *σ* is fixed at 0.13, and the sequencing depth *λ* = 8.25 (see Methods for details on these parameters). GloScope was calculated based on GMM density estimation with latent space representation via the first 10 dimensions of PCA.

We can also compare choices in the data analysis pipeline. For example, GloScope relies on a user-provided choice of latent variable representation of the single-cell data. We compare the choice of PCA versus scVI in a wide range of our simulation settings. The most striking difference is in detection of cell-composition differences, where scVI has much less power in detecting differences between the two phenotypic groups than PCA (Additional File 1: Fig. S2). The latent variable representations given by scVI demonstrates much greater variability between samples than the those of PCA (Additional File 1: Fig. S3), potentially resulting in less power to detect the shared phenotypic differences. On the other hand, scVI representations have more power than their PCA counterparts when the source of differences is due to log-fold changes in genes (Additional File 1: Fig. S4), perhaps due to better accounting for sparse low-count data.

Finally, we can also consider choices made in implementing GloScope, in particular in the choice of estimation of the density of the latent variables *Z* in each sample. We consider two popular density estimation strategies: parametric Gaussian mixture models (GMMs) and non-parametric *k*-nearest neighbors (kNNs). We do not observe large differences in the power of these methods when varying the level of differential expression (Additional File 1:. Fig. S4), but kNN is somewhat more powerful in the presence of cell-type composition changes (Additional File 1: Fig. S2). Applying both methods on a wide range of datasets (Additional File 1: Fig. S5S6) shows that, on average, the estimates of divergence from the two methods are generally monotone with moderate to strong correlations (Pearson coefficient ranging from 0.36 to 0.95); furthermore, the kNN estimates are systematically lower and appear to saturate when GMM estimates are large. While kNN-density estimation offers an asymptotically unbiased estimator of the symmetric KL divergence [25], it is known to exhibit downward finite sample bias due to underestimation of density in the tails of a distribution [26–28]. Due to these considerations, we relied on GMM estimates of density, though none of the results shown qualitatively change if kNN estimates are used instead.

### 2.4 GloScope representation for Quality Control

Finally, we demonstrate the use of GloScope for exploratory data analysis of relatively large sample cohorts and illustrate the utility of having a sample-level representation of the data for exploratory data analysis.

The first dataset is a study of COVID-19 [29] consisting of 143 samples of peripheral blood mononuclear cells (PBMC); samples in the study originated from patients that were either identified as infected with COVID-19 with varying levels of severity (COVID), negative for COVID-19 (Healthy), healthy volunteers with LPS stimulus as a substitute of an acute systemic inflammatory response (LPS), or having other disease phenotypes with similar respiratory symptoms as COVID-19 (non-COVID). Fig. 7A shows these samples after applying MDS to the pairwise divergences calculated from the GloScope representation for the 143 samples of the study.

**Fig. 7.**
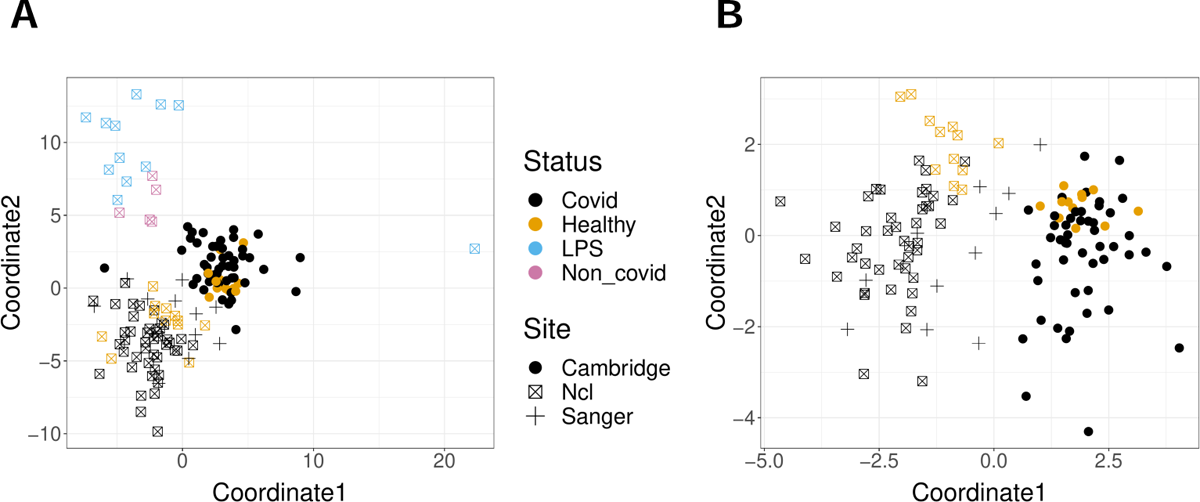
GloScope representation applied to samples sequenced in Stephenson et al. [29]. Shown are the MDS representation in two dimensions of the KL divergence estimates calculated from the GloScope representation for (A) all 143 samples and (B) the subset of 126 samples that were either healthy or diagnosed with COVID-19 (MDS was rerun on the reduced subset of divergences between these 126 samples). Each point corresponds to a sample and is colored by the sample’s phenotype; the plotting symbol of each sample indicates the site at which the sample was sequenced (see legend). Estimated GloScope divergences used the GMM estimate of density and latent variables were estimated with PCA in 10 dimensions. For the visualization of the full divergence matrix, see Additional File 1: Fig. S13.

The visualization shows that both COVID patients and healthy donors are clearly separated from patients with other respiratory conditions (LPS and non-COVID). The other noticeable pattern is that the remaining patients do not show a strong separation between the COVID and Healthy phenotypes, but do appear to separate into at least two groups unrelated to these main phenotypes of interest – an observation that is further strengthened when considering the MDS representation of only the COVID patients and healthy donors (Fig. 7B). Exploration of the provided sample data from Stephenson et al. [29] shows that these groups correspond to different sequencing locations, indicating a strong batch effect due to sequencing site, with samples sequenced at the Cambridge site clearly separated from those at the New Castle (Ncl) and Sanger sites. When the individual cells are visualized (Additional File 1: Fig. S14), the distributional differences between these sequencing sites validate these differences, with cells from the Cambridge site lying in quite different spaces from cells of the same cell type from the other sequencing sites. Furthermore, Stephenson et al. [29] indicates that samples from these different sites underwent different sequencing steps such as cell isolation and library preparations (and the original analysis in Stephenson et al. [29] corrected for potential batch effects by applying the batch correction method, Harmony [10]).

A similar analysis was applied to a Systemic lupus erythematosus (SLE) dataset, with scRNA-Seq data of the PBMC cells of 261 patients; some patients had multiple samples resulting in total 336 samples [30]. Again, our GloScope representation clearly shows that there are distinct patterns among different batch sources, in addition to separation of normal samples from the other conditions (Fig. 8A). After application of Harmony to this data based on the batch, our GloScope representation shows much greater intermingling of the data from different batches (Fig. 8B). We can quantify the improvement by measuring the separation between samples within a batch compared to those in separate batches using measures such as the ANOSIM *R* statistic or Silhouette width. We see the improvement due to batch correction, but some loss of separation between biological conditions, which is a common trade-off when correcting for batch effects (Fig. 8C and 8D). This type of exploratory analyses of data is a common task in the analysis of scRNA-Seq data, and the GloScope representation provides a meaningful strategy for evaluating these types of processing choices. We further note that in addition to finding differences amongst the sequencing sites in the Lupus PBMC data, we observe further clustering of samples in Batch 4 (highlighted in Fig. 8A). These subgroups do not correspond with any patient covariates provided by the authors, but further exploration clearly show strong differences in the gene expression and cell density in certain cell types such as CD4 T cells, Natural Killer cells, and B cells. (Additional File 1: Fig. S16 and S18).

**Fig. 8.**
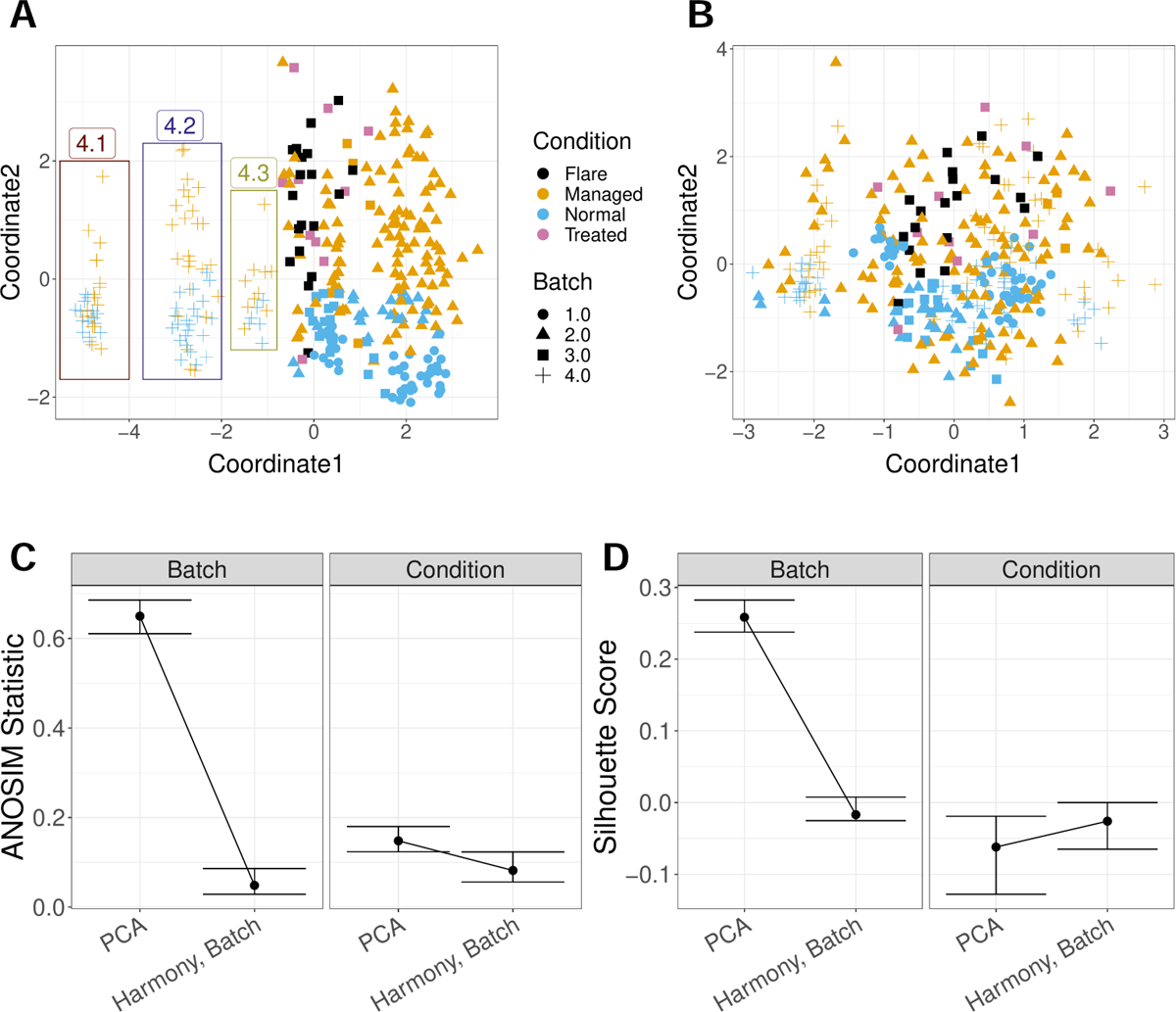
GloScope representation applied to a Systemic lupus erythematosus (SLE) PBMC dataset of 336 samples. [30]. Shown is the MDS of the GloScope representation applied to latent variables defined by (A) the first 10 PCA components of the original data and (B) the latent variables defined by Harmony after normalizing on processing cohort. (C) The ANOSIM statistics changing regarding capturing batch or condition signal, before and after applying batch correction (i.e Harmony) with bootstrap confidence interval. (D) the Silhouette widths changing regarding capturing batch or condition signal, before and after applying batch correction with bootstrap confidence interval.

Similar concerns are frequently explored when integrating data from different studies. We applied GloScope on the dataset of Fabre et al. [31] which integrated six lung fibrosis scRNA-Seq studies, resulting in 144 samples after quality control. Application of GloScope (Fig. 9A) immediately shows one of the studies [32] as quite different from the other five; further investigation shows that the study of Adams et al. [32] has quite obvious differences in both gene expression and cell type composition than the other five studies. In particular, we observed quite obvious gene expression shifting in myeloid cells and Natural Killer cells in Adams et al. [32] (See Additional File 1: Fig. S21), and samples collected from Adams have a higher portion of myeloid cells compared to samples from other studies (Additional File 1: Fig. S20). The remaining five studies show relatively smaller differences, but some separation is clearly visible.

**Fig. 9.**
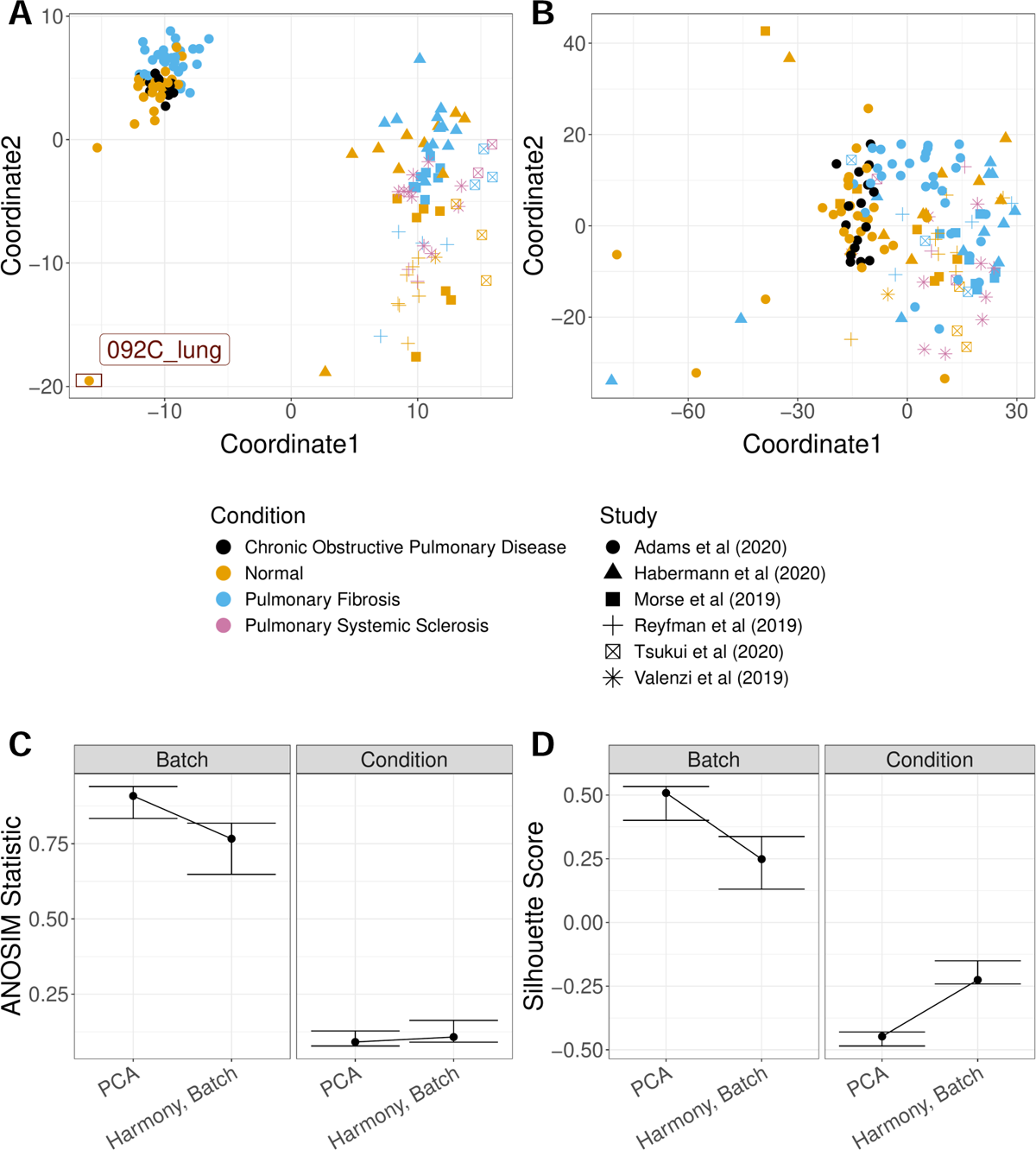
GloScope representation applied to lung fibrosis samples collected in Fabre et al. [31]. Shown are the MDS representation in two dimensions of the KL divergence estimates calculated from the GloScope representation for (A) PCA embedding before batch correction and (B) PCA after applying Harmony batch correction. Each point corresponds to a sample and is colored by the sample’s phenotype; the plotting symbol of each sample indicates the studies at which the sample was collected (see legend). Estimated GloScope divergences used the GMM estimate of density and latent variables were estimated with PCA in 10 dimensions. (C) and (D) visualize the ANOSIM R statistics and Silhouette width, quantifying the changes of batch and biological signals before and after batch correction

In addition to large batch effects, we observed a potential outlier (sample 092C lung), from the Adams et al. [32] study detected by the GloScope representation (Fig. 9A). Further evaluation of that outlier sample shows that 092C lung is missing most of the cell types except for B cells and lymphocytes (Additional File 1: Fig. S22). In contrast, a similar analysis of data from [31] which integrated studies of 50 human liver samples (after quality control) from 6 published scRNA-Seq studies of liver fibrosis shows far less distinction among the studies compared to the lung samples (Additional File 1: Fig. S23). Following application of Harmony for batch correction/integration, the GloScope shows effective integration of the lung studies and a corresponding clearer grouping of biological conditions. (Fig. 9C,D).

Batch effects are common concerns with large sets of data, especially in human subject data where the samples are likely to be collected and possibly sequenced at different sites or integrated across multiple smaller studies. These examples immediately demonstrate the power of our GloScope representation for exploratory data analysis.

### 2.5 Comparison with other Quality-control tools

Existing tools for EDA and evaluation of potential quality concerns are generally focused on analysis at the level of the individual cell. Numerous metrics exist for evaluating the quality of individual cells and filtering poor cells, such as the the number of detected genes, the number of sequenced reads, or the percentage of mitochondrial DNA [33, 34]. Yet, many sources of possible artifacts are often due to variables that vary per sample or patient, such as the hospital of collection, the sequencing site, or the laboratory running the experiment. These effects have large-scale effects beyond individual cells and are best detected by comparisons of the cells as a group. However, there are limited options for detecting artifacts that vary by sample or individually poor samples.

In particular, analyses at the individual cell-level are less flexible for detecting these sample-level differences. There are metrics at the individual cell-level, such as iLISI [10] that can assess the presence of a batch effect *for known batch variables*. These are similar to our use of ANOSIM or Silhouette width to quantify the separation between samples in batches, only these methods are applied to the individual cells. Such methods can highlight similar effects, such as showing an improvement in Harmony corrected data for the Stephenson et al. [29] data (Additional File 1: Fig. S24), but they are ineffective for discovering effects *de novo*, nor do they provide the ability to compare multiple effects, such as our visualizations of both batch and biological effects in Section 2.4.

A common exploratory visualization strategy for scRNA-Seq data consists of applying tools such as UMAP or tSNE to create a two-dimensional visualization of the individual cells. Individual cells can be color-coded by potential variables or plotted separately per sample for exploration of possible *known* artifacts, as we provided for the [29] data in Additional File 1: Fig. S14. UMAP visualizations can be helpful in retrospect for understanding the nature of the problem, but are not particularly effective in discovering such effects *de novo* given the difficulty in visualizing sample effects for large numbers of cells. The example of the Perez et al. [30] data is illustrative, where our GloScope representation allowed us to immediately determine unexplained groupings of samples within Batch 4; we were able to follow this discovery with further investigation at the individual cell-level using UMAPs to discover that there were shifts in gene expression and cell density among these subgroups GloScope identified within Batch 4. These differences are not detectable in plotting all cells, and only after identifying the subgroups of patients can a UMAP help in further investigation. Furthermore, differences due to shifts in cell distributions can be tricky to see in UMAP visualizations of individual cells, due to the overplotting of cells. Even after identifying the different subgroups in Batch 4 with GloScope (Fig. 8), the differences seen clearly in the GloScope representation were subtle to detect using standard UMAP visualization (Additional File 1: Fig. S18, S17, and S19). This exploratory analysis of the [30] data shows the complementary nature of GloScope with other visualization tools. Similarly, outlying individual patients, as we detected in the lung samples of Fabre et al. [31] (Section 2.4), would require plotting and comparing of UMAPs of each individual sample which is simply not feasible for large cohorts.

There are some limited alternatives to GloScope available for the comparison at the sample-level, and they take different strategies for summarizing the data from a single patient which we next consider: cell-composition and pseudobulk.

### Comparison with cell-composition analysis

Reducing each sample to their cell-type composition has been proposed for comparing samples. A simple version of this strategy is to visualize the proportions per sample in a barplot. Like UMAPs of individual cells, such barplots can be useful tools for greater investigation of differences found by GloScope, but do not scale for easy comparisons of large number of samples and do not aid in discovering possible differences, such as the potential subgroups of batch identified by GloScope (Additional File 1: Fig. S16, S15).

The cell-type proportions can also be analyzed more quantitatively– for example the GloScope methodology can also be used for cluster proportions, which we call GloProp (see Section 2.1). Concurrently, Joodaki et al. [16] has proposed a similar metric strategy for comparing cell-type proportions named PILOT, using Wasserstein distance rather than symmetric KL divergence. These approaches require determination of cell-type proportions and can only be run after clustering the individual cells. Such clustering is typically done after EDA and correction of possible batch effects, making it irrelevant for EDA. But in principle clustering could be done earlier in the pipeline for the sole purpose of using PILOT for EDA (the discovered clusters would not be biologically meaningful until the data has been appropriately pre-processed).

We do this clustering on the uncorrected data and compare PILOT and GloProp to GloScope. We see that PILOT performs much worse than GloScope or GloProp in detecting separations between the batches in all of the datasets (Additional File 1: Fig. S25). Our method for cluster proportions, GloProp, can perform similarly to that of the full GloScope representation, but the performance of both PILOT and GloProp is very sensitive to the clustering. When we vary parameters of the clustering algorithm or consider different random starts, the performance can vary dramatically, unlike GloScope (Additional File 1: Fig. S26).

### Comparison with pseudo-bulk analysis

Another potential strategy for sample-level exploratory analysis is using a pseudobulk created from the scRNA-Seq data. This is a strategy of aggregating over each sample’s cells to obtain a single observation per sample [1]; the most common is to simply sum the counts. Then standard methods from bulk mRNA-Seq, such as PCA, can be applied at the sample level. The authors of [35] propose a strategy, MOFA, for finding lower-dimensional latent embeddings per sample based on combining pseudobulk measures per cell-type, to better reflect cell-type variability.

We create such a PCA visualization of the pseudo-bulk of several of the datasets mentioned above (Additional File 1: Fig S27, S28). For the COVID-19 PMBC samples, for example, the pseudobulk analysis does not clearly separate out the LPS and non-COVID samples, nor is the strong batch effect due to sequencing site as clearly identified. Similarly, for the Lupus PBMC data, the pseudobulk representation does not identify the strong batch effects seen in our GloScope representation. This is borne out by the quantification of the average silhouette width or *R* statistic (Additional File 1: Fig. S29 and S30). On the other hand, these quantification statistics show MOFA to have similar performance in detecting batches as GloScope; however, on closer examination of the visualization of the results of MOFA, we see less clear separation of the effects seen by GloScope. For example, MOFA did not show clear of a separation of all the non-COVID and LPS samples from other samples and the separation of the groupings found *de novo* by GloScope are attenuated and difficult to find (Additional File 1: Fig. S28).

There are other limitations to either of these pseudo-bulk strategies. The pseudobulk strategy, including MOFA, is based on summarizing for each gene the expression level of all the cells in a sample, usually the sum of the raw counts. However, in many public datasets provide other normalized versions of the data (e.g. residuals); similarly many batch-correction methods, like Harmony [10], provide a batch-corrected latent variable representation. None of these are obvious candidates for either of these pseudo-bulk approaches. Our GloScope representation requires as input only a latent-variable representation per cell and thus is flexible to accommodate all of these types of input. This is important, for example, in evaluating the effect of batch correction methods. With GloScope, we can evaluate the data before and after batch correction with the Harmony algorithm (Fig. 8B,C,D), allowing us to confirm that the Harmony algorithm has removed much of the differences between batches. Moreover, the pseudobulk methods can often need normalization across samples in addition to normalization that may be done to individual cells so that they do not reflect simply the number of cells, similar to bulk RNA, which adds another layer of complexity since there are many strategies for such a normalization. GloScope summarizes the individual cells as a density, which is a measurement unaffected by the number of cells per sample.

## 3 Discussion

In this work, we demonstrated the use of GloScope for exploratory analysis, and in particular how the GloScope divergences can be used to create two-dimensional scatter plots of samples, similar to that of PCA plots of bulk mRNA-Seq data. We demonstrated the ability of the GloScope representation to detect important artifacts in the data, as well as assess batch-correction methodologies.

We also compared GloScope to the limited available strategies for summarizing the data from a single patient: cell-type composition and pseudobulk. We show that these methods are not as sensitive in as diverse of settings. In particular, these approaches each focus on one aspect of the sample data (cell-type proportions or gene expression) and are not sensitive to changes found in the other. GloScope uses the entire distribution of the data, thus effectively combining both cell-type proportions and gene expression in a single summary. Furthermore, GloScope is far more flexible for incorporation at different stages of the analysis, whether working with raw counts or normalized data.

While we focus on the utility of the GloScope representation to visualize scRNASeq data at the sample level, the representation can be used more broadly with other statistical learning tools. For example, we can use the GloScope divergences between samples as input to a prediction algorithm in order to predict a phenotype. With the COVID-19 data, we apply the SVM algorithm to the GloScope divergences which results in a prediction algorithm that was able to separate the normal from the COVID samples with a 5-fold cross-validated prediction accuracy of around 0.88. This simple example serves as an illustration of the power of a global representation of the entire scRNA-Seq profile.

Finally, we note that GloScope can easily be incorporated into existing scRNA-Seq pipelines at multiple stages of analysis to assess the progress. Latent-varible representation, via PCA or scVI is a standard initial step in an analysis, while many popular batch correction methods provide low-dimensional representations of corrected data. Even multi-modal integrations usually result in a low-dimensional latent space estimation. The output of all of these tasks can be provided to GloScope for evaluation of sample-level similarities, resulting in a flexible tool for exploratory analysis of the results.

## 4 Conclusion

We have presented the statistical framework GloScope that provides a global summary of each scRNA-Seq sample based on the distribution of their gene expression values across their cells. This representation allows for comparisons between the entire singlecell profile of a sample. Formal calculations of the dissimilarities between samples can be used as input to other statistical and machine learning algorithms to allow a sample-level analysis. Our representation is able to differentiate among samples from varied phenotype groups, such as COVID lung tissue samples and healthy lung tissue samples, and is shown to be a powerful tool to detect potential batch effects.

## 5 Methods

### 5.1 The GloScope representation

Our GloScope representation consists of representing each sample as a distribution along with a corresponding divergence or distance; we then estimate the distance or divergence between each pair of samples based on their scRNA-Seq data. This representation allows for application of kernel methods common in machine learning, which depend on the calculation of the distance between each pair of samples *i, j* for downstream statistical analysis.

To do this, we posit an underlying true distribution of cells *F_i_* for each sample *i*, which is a continuous probability distribution on *R^g^*, where *g* is the number of genes. We define a measure of divergence *d* on the space of probability distributions in *R^g^*. In this work, we fix *d* as the symmetrized Kullback-Leibler divergence,

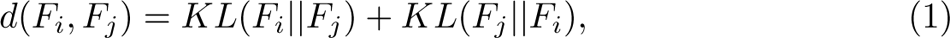

which has been used in a similar manner in the case of facial recognition [e.g., 6, 36, 37]

We do not observe the *F_i_* directly and must instead estimate that distribution from observed data. The observations from a sample *i* consists of *m_i_* sequenced cells; in order to estimate *F_i_* we will make the simplifying assumption that the sequenced cells are independent and identically distributed (i.i.d) draws from the sample’s full population of cells, *F_i_*. Even with this assumption, density estimation is complicated in this setting. For scRNA-Seq datasets, *g* is often in the range of 2,000-8,000 (the number of detectable genes given the sequencing depth). The number of cells per sample, *m_i_*, can vary by experiment, and often *m_i_* ranges lies in the range of 500 to 10,000 cells per sample. The data from each cell is high dimensional and sparse, a distributional structure known to be impactful in the analysis of scRNA-Seq data [38–42].

### Defining a Latent Space

Even with several thousand cells per sample, it is infeasible to estimate the density in such a high dimensional space without the assumption of an underlying lower dimensional latent space. Therefore, for each sample *i* and cell *c* we model a latent variable *Z_ic_ ∈ R^d^* and a transformation *σ*: *R^d^ → R^g^*. Then our observed vector *x_ic_* of gene expression counts from a cell is assumed drawn from an appropriate generative model for RNA counts with mean parameter *σ*(*Z*), i.e. *E*(*x_ic_*) = *σ*(*Z_ic_*).

For a sample *i*, we assume that the *Z_ic_* for each cell *c* is distributed as the latent random variable *H_i_*. Instead of estimating *F_i_* in *R^g^*, GloScope instead estimates *H_i_* in the lower-dimensional space *R^d^*. In Section 2.1, we denote the estimated distribution as *F*^^^*_i_* for conceptual simplicity, but a more precise notation would be *H*^^^*_i_* to clearly emphasize that we are estimating the distribution on a lower-dimensional space.

Furthermore, we note that our above heuristic states that we observe counts *x_ic_* in cell *c* drawn from a single distribution *F_i_*; this ignores cell-specific effects that could result in slightly different distributions for different cells, such as different sequencing depth that varies for each cell *c*. The latent variables *Z_ic_*, however, are independent of the cell-specific effects due to the technology, which makes estimation of a single distribution, *H_i_*, shared by all cells a coherent mathematical framework.

### Estimation

The GloScope representation estimates *H_i_* for each sample with a two-stage strategy: 1) estimation of the latent variables *Z_ic_ ∈ R^d^* for each cell *c* in sample *i* and 2) estimation of the density of *H*^^^*_i_* from *Z_ic_* and corresponding distances *d*(*H*^^^*_i_, H*^^^*_j_*) between samples. An advantage of estimating the latent variable samples before the density is that we can apply one of many existing dimensionality reduction techniques that account for sparse count data, such as ZINBWave [39] or scVI [5], or techniques that simultaneously remove batch effects and estimate a latent space, such as Harmony [10] or fastMNN [43].

The GloScope representation assumes that the user chooses an appropriate method for the first stage estimation of *Z_ic_* (i.e. a dimensionality reduction method) and then offers two approaches for the second stage (estimation of the distances between the *H_i_*).

The first approach applies a Gaussian mixture model to the *Z_ic_* to estimate *h_i_*, the density associated with the distribution *H_i_*, and then calculates *d*(*H*^^^*_i_, H*^^^*_j_*) as our estimate of *d*(*F_i_, F_j_*). Single cell methods utilizing dimensionality reduction, described above, often include a regularizing assumption that the latent variables *Z ∼ N* (0, Σ). This Gaussian regularization in the model and the fact that many datasets are mixtures of cell type populations, motivates our use of Gaussian mixture models (GMMs). We use the R package mclust [44] to implement the GMM estimation. As there is no closed form expression for the KL divergence between GMM distributions, we use Monte Carlo integration to approximate the KL divergence between two GMM densities; this is based on *R* = 10, 000 samples drawn from the estimated GMM distributions, again using the mclust package. Specifically, for *R* draws of *x* from *H*^^^*_i_*, we have

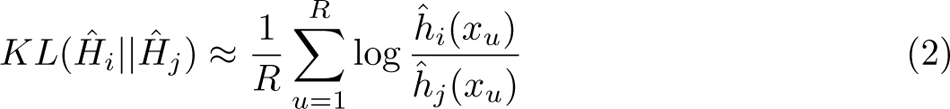

We also provide a second approach that estimates *d*(*H_i_, H_j_*) directly using a knearest neighbor approach without explicitly estimating the density *h_i_* [45, 46]. Denote by *r_j_*(*x_i,u_*) the distance from the *u*th cell in sample *i* to its kth nearest neighbor in sample *j*. Then the KL divergence can be estimated directly as

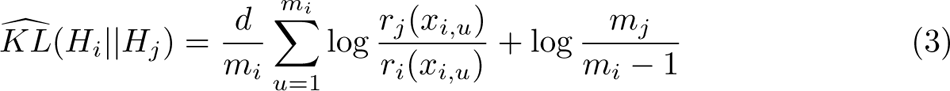

where d is the dimension of the latent space [45, 46]. We implement this strategy using the FNN package to estimate the symmetrized KL divergence between sample *i* and sample *j* [47].

### 5.2 Simulating scRNA-Seq data

#### 5.2.1 Simulation Model

To simulate population-level scRNA-Seq data with which we benchmark our methodology, we follow the model introduced by the muscat R package. We would note that this is a model for simulating count data for each gene, and unlike our GloScope representation does not assume any latent variable representation in generating the data. The muscat package assumes a simple two-group setting in which each sample *i* may come from one of two groups, denoted by the variable *T* (*i*) *∈ {*1, 2*}*. The *m_i_* cells from sample *i* come from *K* different cell-types with the proportion of cells from cell-type *k* given by *π_i,k_*, where ^L^*_k_ π_i,k_* = 1. Thus the gene expression vector *x ∈ R^g^* of a cell *c* from sample *i* is assumed to follow a negative binomial mixture model:

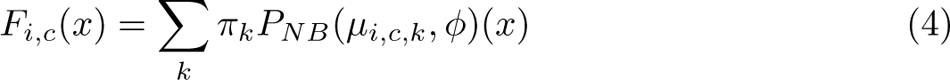

where *P_NB_*(*µ_i,c,k_, ϕ*) is a CDF on *R^g^* representing a product distribution of independent negative binomials, i.e. each gene’s expression value is independent and follows a negative binomial distribution with mean given by the *j* the element of the vector *µ_i,c,k_ ∈ R^g^* and dispersion parameter *ϕ ∈ R*.

The vector of gene means for cell *c* in sample *i* is parameterized in `muscat` as

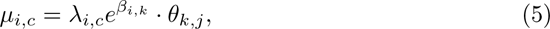

where *λ_i,c_ ∈ R* is the library size (total number of counts); *β_i,k_ ∈ R^g^* is the relative abundance of *g* genes in cells belonging to sample *i* and cell-type *k*; *θ_k,j_ ∈ R^g^* is the fold-change for genes in cluster k if the sample belongs to group *j ∈ {*1, 2*}*. Notice, as mentioned above, that because of different sequencing depths per cell, each cell within sample *i* has a different mean *µ_i,c,k_* governed by the sequencing-depth parameter *λ_i,c_*, hence our notation *F_i,c_*.

We make adjustments to the above model in the muscat package to more fully explore sample variability. To explore the effect of library size variation at both the cell and sample level, we introduce the decomposition *λ_i,c_* = *λ*^-^ + *λ_i_* + *δ_c_,* where *λ*^-^ is the overall (average) library size, and *λ_i_* and *δ_c_* are variations from that due to sample or cell level differences, constrained so that *λ_i,c_ >* 0. We also adjusted the model to allow sample-specific proportions vectors *π_i,k_*, with ^L^*_k_ π_i,k_* = 1. We define proportions per treatment group, Π*_j_ ∈ R^K^*, for treatments *j* = 1, 2, such that ^L^*_k_* Π*_j,k_* = 1 and randomly generate probability vectors *π_i_* for sample *i* from a Dirichlet distribution according to its treatment group, *π_i_ ∼ Dirichlet*(Π*_T_* _(_*_i_*_)_*∗α*), with sample level variation parameter *α*.

### Selection of Parameters

The muscat package also provides methods for creating these many parameters based on a few input parameters by the user and estimating the other parameters based on reference data provided by the user. We followed their strategy, with the following additions.

We chose the group fold change difference per cell-type, *θ_k,j_* following the schema of muscat, which allows for various types and size of changes between the different groups. Briefly, the simulation of *θ_k,j_* is controlled by parameters 1) Ω *∈ R*, which is a user-defined average log2 fold change across all DE genes, 2) *ω_k_ ∈ R^k^*, which varies the magnitude of gene expression difference for cluster k, and 3) a proportion vector *ρ* which is the proportion of genes that follow six different gene expression patterns (see [1]); for simplicity, we allowed only the two most typical gene expression patterns, which are EE (equally expressed) and DE (differentialy expressed) genes for our simulations, resulting in *ρ* effectively being a single scalar, the proportion of genes that are differentially expressed.

The selection of *m_i_*, the number of cells per sample *i*, also followed the strategy of muscat, where the user provides a value *m*^-^, representing the average number of cells per sample across all samples, and the value of each individual *m_i_*for each sample is assigned via a multinomial with equal probability and total number of cells across all samples equal to *n ∗ m*^-^.

The parameters *ϕ*, and initial values of *λ_i,c_*and *β_i,k_* were obtained by estimating these parameters from the reference data, following the muscat package: after performing quality control, we used the filtered gene matrix and the edgeR package to estimate the parameters from the reference data.

Using our modified parameterization described above, *λ*^-^ was then chosen as the average of the *λ_i,c_* estimated from the reference samples. Sample-level sequencing depth variability *λ_i_* were simulated as *λ_i_ ∼ Unif* (*−τ_λ_, τ_λ_*). Per-cell variability, *δ_c_*, was simulated as *δ_c_ ∼ Unif* (*−τ_δ_, τ_δ_*).

Finally, the selection of *β_i,k_* used in our simulation diverged from muscat package strategy. The muscat estimates of *β_i,k_* created overly large differences between the treatment groups and samples (Additional File 1: Fig. S11); furthermore their strategy recycles the same set of parameters *β_i,k_* if the simulated sample sizes are larger than provided reference sample sizes (i.e. the same value of *β_i,k_* would be given to multiple simulated samples), resulting in unintended batches of samples. Instead, we estimated *β*^^^*_i,k_* from the reference data using the muscat strategy, and chose a single sample *i^∗^* whose initial estimates *β*^^^*_i,k_* were representative. We then set *β*^^^*_k_*= *β*^^^*_i_∗_,k_* and created individual *β_i,k_* with variation per sample by adding noise to *β*^^^*_k_*, *β_i,k_* = *β*^^^*_k_ /*2 + *ξ_i,k_*, where *ξ_i,k_ ∼ N* (0*, σ_ξ_*). *σ_ξ_* controled the degree of sample-level variation.

Additional File 1: Fig. S12 shows the effect of changing different parameters (*σ* and log-fold change), visualized using UMAP on an illustrative example.

#### 5.2.2 Simulation Settings

In following the above strategy of selecting parameters, we randomly chosen 5 COVID samples from the COVID-19 PBMC dataset, [29]. After estimating *ϕ* and *β*^^^*_k_* as described above from the reference samples, the values were fixed for all simulations.

The value *m*^-^ was chosen as 5,000, which is similar to the average cell per samples in several datasets (e.g. [22, 23, 29]). The default value for *α* to control the sample level cluster proportion variability was set to be 100, except where explicitly noted, which keeps the variation in cluster proportions to be relatively small among samples (see Fig. 5D).

Once these parameters were fixed, the following user-defined parameters were set differently for different simulation settings: *n* (the number of samples in a single group), the vector group proportions Π*_j_* (*j* = 1, 2), average library size *λ*^-^, and the DE parameters Ω, *ω*, and *ρ*. With these global parameters chosen for a simulation setting, the remaining sample-specific parameters are generated anew in each simulation:

1. for each cell-type *k*, *n* values of *β_i,k_* as described above based on *β*^^^*_k_*,
2. for each cell-type *k*, a single vector *θ_k,j_ ∈ R^G^* for the population log-fold-change between groups, based on the parameters Ω, *ω*, and *ρ*,
3. for each sample *i* a single value *λ_i_* and *m_i_* values of *δ_c_*, one for each of the *m_i_* cells from each sample. This results in *m_i_* values of *λ_i,c_* = *λ*^-^ + *λ_i_* + *δ_c_* for each sample. (Note that some simulations set *λ_i_* and/or *δ_c_* to 0 for all *c* and *i*).

Combining these parameters result in the *µ_i,c,k_* needed for each sample in a single simulation, and then the cell-counts for each sample *i* are simulated from *F_i,c_*. Additional Files 2-7 provide the different parameter settings that were run and their resulting power and average ANOSIM values corresponding to the figures shown here.

#### 5.2.3 Numerical metrics for evaluating simulations

In order to quantify how well our representation was able to differentiate sample groups in different settings, we implemented a simple hypothesis test for comparing the two groups based on our estimated distances from our GloScope representation. We relied on the Analysis of Similarities (ANOSIM) test, which is a non-parametric test based on a metric of dissimilarity, to evaluate whether the between group distance is greater than the within group distance. We used the function *anosim* in the R package vegan to perform the test [24]. The test statistic is calculated as:

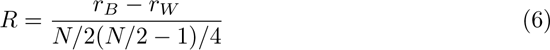

where *r_B_* is the mean of rank similarities of pairs of samples from different groups, *r_W_* is the mean of rank similarity of pairs within the same groups, and *N* is the total number of samples. The test statistics ranges from −1 to 1. Strong positive test statistics means greater between group distances than the within groups; strong negative test statistics means the opposite and may represent wrong group assignments; and test statistics near zero indicate no differences. Finally, p-values are calculated based on a null permutation distribution: the distribution of *R* recalculated after randomly shuffling the samples’ group assignment. The p values are calculated as the proportion of times that the permuted-derived statistics are larger than the original test statistic. We used the results of ANOSIM to calculate the power in different simulation settings, creating a quantitative metric for evaluating the sensitivity of the GloScope representation in different scenarios. For a choice of input parameters, we repeated the simulation 100 times. For each simulation, we calculated the pairwise distances between all 2*n* samples, then used ANOSIM p-values to determined whether we would reject the null hypothesis. Finally, we calculated the power as the proportion of the 100 simulations’ test statistics that have p values smaller than *α* = 0.05.

### 5.3 Data processing procedures

This section details the steps undertaken to estimate GloScope representations of samples from publicly available scRNA-Seq data. These steps broadly consisted of ensuring the data we used had quality control matching the corresponding paper, estimating the cells’ latent embeddings, and applying the GloScope methodology. For most datasets we performed the first two steps with data structures and functions from the R package `Seurat`. For the larger Lupus PBMC and mouse brain datasets, we instead utilized the `SingleCellExperiment` data structure and applied functions from other packages. Code for running these analyses, as well as text files containing data sources and specific processing choices, are available in the following GitHub repository: https://github.com/epurdom/GloScopeanalysis [48].

#### Quality Control Verification

The UMI count data and cell annotations from each sample-level scRNA-Seq study were downloaded from its publicly accessible source (indicated in the code). We checked whether data provided already had the quality control steps described in its respective paper. These steps can include removing cells with extreme expression values and filtering certain gene sets, such as mitochondrial genes. Only the data provided from Ledergor et al. [49] did not appear to have the stated steps of the manuscript already applied, and we reproduced the cell-wise quality control procedure described in that paper’s Methods section. We also removed genes expressed in less than 10 cells (except for the data from Stephenson et al. [29] which provided only PCA embeddings that we used directly, and Fabre et al. [31], where the preprocessed and normalized data is provided).

#### Latent Space Estimation

In this paper we present results based on using 10-dimensional latent embeddings, calculated with either scVI or PCA. To calculate scVI embeddings, we used the entire UMI count matrix as input after the aforementioned verification steps. To calculate PCA embeddings, we used a subset of only the 2, 000 most highly variable genes. To select which genes to include, we first log-normalized the counts within each cell; this was implemented with *logNormCounts* from Seurat or *logNormCounts* from `scuttle` for `SingleCellExperiment` objects. Then we fit a LOESS curve to predict each gene’s log-scale variance from its log-scale mean; that regression was implemented with the *vst* method of *FindVariableFeatures* in Seurat and with *modelGeneVar* from the scran package for `SingleCellExperiment` objects. The *FindVariableFeatures* function of Seurat also selects the 2, 000 highly variable genes based on large residuals in the LOESS regression. That exact selection rule is not available for `SingleCellExperiment` objects, so we instead applied a similar procedure implemented by *getTopHVGs* from scran. This alternative only differs in a truncation step and is commonly used in other scRNA-Seq analyses [50]. Each of the 2, 000 selected genes was centered and scaled to zero mean and unit variance before running PCA. In Seurat objects this was done via a two-step procedure with calls to *ScaleData* and *RunPCA*. However for `SingleCellExperiment` objects we standardized each gene and ran PCA with a single call to *runPCA* from scater.

#### Application of GloScope

After obtaining each cell’s latent representation via PCA or scVI, we fit sample-level densities with GMM or kNN and the KL divergence between samples was estimated. This produces the GloScope representations, and these steps are implemented by *gloscope* function in our GloScope R package, which accompanies this paper and is available in the accompanying Bioconductor package GloScope (https://www.bioconductor.org/packages/release/bioc/html/GloScope.html, version 1.2.0 [51].

To run GloScope, we first had to determined which cells constituted a single sample in each dataset, based on the provided metadata. In some studies each patient only provided one sample, and in others a single patient provided multiple samples; for instance from affected and healthy regions. Considering this, we ran GloScope with tissue samples as the unit of analysis. The sole exception to this choice is the PBMC data from Lupus patients in Perez et al. [30]; this study processed some tissue samples in multiple processing cohorts, and we therefore used the cross of sample and cohort as our unit of analysis.

Before applying GloScope we confirmed that the tissue sample identifier associated with each cell matches the reported study design. The original melanoma data from Jerby-Arnon et al. [52] had duplicate encodings, which we standardized, and one sample with a mislabeled phenotype. For multiple myeloma cells from Ledergor et al. [49], we parsed concatenated strings into patient, observation period, and phenotype indicators. Two colorectal tumor samples from Pelka et al. [23] were sequenced with two technologies, and we only considered the replicates using the newer technology.

For datasets other than Fabre et al. [31], we also chose to remove samples with less than 50 cells. This excluded 2 samples from Ledergor et al. [49] (AB3178 and AB3195) and 1 from Pelka et al. [23] (C119N). For data in Fabre et al. [31], we observed extremely small cell number per sample (minimum 2 cells) and chose to remove samples with less than 500 cells, resulting in removing 4 samples in Lung study (137CO lung, 244C lung, 8CO lung, and 084C lung), and 15 samples in Liver study (P13 healthy liver, P4 Tumor liver, P8 healthy liver, P10 Tumor liver, P11 Tumor liver, P11 healthy liver, P5 healthy liver, P7 healthy liver, HN healthy liver, P12 healthy liver,P1 healthy liver,P3 healthy liver and P9 healthy liver) We noted that one sample from Ledergor et al. [49] (AB3461), had extreme divergences with other samples. We removed all cells from this sample for the results presented in this paper.

#### Computational Time

In Additional File 1: Table S1 we provide the timing for how long it took to run to GloScope compared with other steps in the pipeline. We would note that there are two major computational tasks: the estimation of the density per sample and the estimation of the divergence between pairs of samples. However, because the calculation of the densities is more time intensive than the estimation of the pairwise divergences, the total time to run GloScope is roughly linear in the number of samples, not quadratic for the sample sizes reasonably encountered in scRNA-Seq studies.

### 5.4 Pseudobulk Comparison Experiments

For datasets where raw count data are available, we performed pseudobulk analysis by summing each sample’s cell entries across each genes using *rowSums*, a base R function. After obtaining the pseudobulk data, we processed it using functions from the Seurat package. We log-normalized the data using *NormalizeData* and selected the top 2000 highly variable feature using the *FindVariableFeatures* function with default arguments. Counts from the selected genes were scaled using *ScaleData* and a PCA embedding was obtained with *RunPCA*.

To perform MOFA analysis, we followed the vignettes provided in the R package MOFAcellulaR [35]. For filtering cells for downstream analysis, we set *ncells* to be 1. As we used the pre-processed data, we did not filter on sample and set *nsamples* threshold to be 0. For the final output, we set *num factors* to be 5.

### 5.5 Comparison to Cluster Composition

In Section 2.5, we compare our methods to those that summarize the samples using cluster composition. We call it “cluster composition” rather than “celltype composition” to emphasize that this is clustering done before standard batch corrections, and thus the results are not biologically meaningful. To do this comparison, we calculate cluster assignments for the individual cells based on a variety of different clustering choices: different algorithms and various choices of parameter settings and random starts for those algorithms. For each clustering assignment generated, we calculate the proportion of cells that are classified into each cluster, and use those cluster proportions as input to PILOT [16] and GloProp, our implementation of GloScope for proportions.

To generate cluster assignments for individual cells, *de novo* clustering is preformed on lower-dimensional cell representations. In our implementation, we focus on PCA to match our input into GloScope. We use two of the most utilized clustering algorithms in scRNA-Seq for this task: the Louvain algorithm [53] and its refinement in the Leiden algorithm [54]. Leiden algorithm is recommended by recent review studies [55, 56] and has become the default clustering algorithm for popular software tools such as scanpy [57] and Monocle3 [58]; furthermore, the Louvain algorithm codebase is no longer maintained. We present only results from Leiden, but the results from Louvain were qualitatively similar.

Both algorithms expect a *k*-nearest neighbor graph between cells as the input.

In these experiments we present results utilizing a *k*-nearest neighbor graph with *k* = 20, the default in the Seurat package [59]. Clustering result initalized by graphs with *k* = 5 and *k* = 100 were qualitatively similar. Both algorithms also have a resolution parameter as the primary parameter, a value which is correlated with the number of clusters ultimately identified; larger values of this parameter lead to fewer unique clusters. For the Leiden algorithm implemented with igraph [60], the default resolution parameter is 1, which is also the default in scanpy and similar to the default of 0.8 in Seurat; however, this value led to each cell being placed in its own cluster.

Instead, we followed the example of the *cluster cells* function in Monocle3 and set our default resolution parameter to be 10*^−^*^5^; this parameter choice gave rise to a similar number of clusters as the default parameter of the Louvain algorithm as implemented in Seurat. We also repeated this experiment with resolution values in *{*10*^−^*^4^, 5*^−^*^5^, 5*^−^*^6^*}* (Additional File 1: Fig. S26); these values were selected to give an average number of unique clusters ranging from tens to hundreds, a representative range of cell types identified in single-cell studies. For each parameter value, we considered results from 20 different random seeds.

### 5.6 Numerical metrics for evaluating performance

To evaluate different methods’ performance on detecting batch effects or biological signals, we used the two metrics for quantification:

1. ANOSIM *R* statistic, equation 6 above,
2. Average Silhouette width using *silhouette* in R package cluster.

After obtaining the above two values from the calculated distance matrix *D*, we calculated bootstrap confidence intervals for each of the metrics. To do so, we defined the unique combinations of batch and biological condition. For each unique combination, we repeatedly sampled with replacement from samples in that combination; the union of the sampled samples from each combination resulted in a single full bootstrap sample. After obtaining the bootstrap sample for each run, we obtained the bootstrap distance matrix *D_boot_* from the original distance matrix *D* by subsetting to the bootstrap sample ids. Finally, we calculate the two metrics based on *D_boot_*. We repeated this for *B* = 100 bootstrap samples. For each of the metrics, we calculated percentile bootstrap confidence intervals by taking the 2.5% and 97.5% quantiles from the empirical distribution of the bootstrap distribution of the metrics.

### 5.7 Prediction of Phenotype with GloScope Representations

After using GloScope to obtain the symmetrized KL divergence matrices of COVID PBMC samples [29], we obtained their MDS embeddings with 10 dimensions. 60% of the data points were reserved for training and rest 40% were used for testing purpose. We applied SVM to classify sample’s phenotype using the package e1071 (i.e, COVID vs healthy), and 5-fold cross valiadtion to tune the hyperparameters cost and *γ* [61, 62]. Finally, we used the test sets to assess the prediction algorithm by counting the prediction accuracy rate.

## Supplementary information

Additional File 1: Supplementary table and figures. Contains supplementary table S1 and figures S1–S30.

Additional File 2: baseline.csv, table for power and ANOSIM statistics calculated based on 100 simulation to detect between group difference when changing gene level variability *β_k_*.

Additional File 3: cluster.csv, table for power and ANOSIM statistics calculated based on 100 simulation, to detect between group difference when changing cluster proportion only.

Additional File 4: lfc pde.csv, table for power and ANOSIM statistics calculated based on 100 simulation, to detect between group difference when changing log-fold change and percentage of DE genes.

Additional File 5: omega k.csv, table for power and ANOSIM statistics calculated based on 100 simulation, to detect between group difference when changing log-fold change concentrated in certain clusters.

Additional File 6: offset.csv, table for power and ANOSIM statistics calculated based on 100 simulation, to detect between group difference when changing library sizes *λ*.

Additional File 7: S power.csv, table for power and ANOSIM statistics calculated based on 100 simulation, to detect between group difference when changing sample size n.

## Acknowledgements

None.

## Declarations

- Funding

This work has been supported by NIH grant 1R01GM144493, NIH grant 5R01DC007235, NSF training grant DMS RTG 1745640, and a Chan Zuckerberg Initative Data Insights Award. EP is a Chan Zuckerberg Biohub investigator.

- Competing interests

The authors declare no competing interests.

- Ethics approval and consent to participate Not applicable.
- Consent for publication

Not applicable.

- Availability of data and materials

Code for running these analyses, generating figures, and text files detailing dataset download source and specific processing choices are available in the following GitHub repository: https://github.com/epurdom/GloScopeanalysis [48]. The GloScope implementation is available in an accompanying Bioconductor package GloScope (https://www.bioconductor.org/packages/release/bioc/html/GloScope.html) [51], with all analyses performed on version 1.2.0.

- Author contribution

HW, WT, BG and EP contributed to the development and modeling work described in the manuscript. HW, WT and EP wrote the main manuscript text. HW and WT developed figures and/or tables for the manuscript. All authors reviewed the manuscript and provided critical editing to main manuscript text. The authors read and approved the final manuscript.

