## Additional File 1: Supplemental Figures and Tables for "Visualizing scRNA-Seq Data at Population Scale with GloScope"

### Supplementary Table

| Metadata | | | Processing | | | GloScope | |
|---|---|---|---|---|---|---|---|
| **Name** | **Samples** | **Avg. Cells** | **PCA** | **Cluster** | **UMAP** | **Total** | **Per Sample** |
| Rash Epidermis [1] | 12 | 7,741 | 0.3 | 0.6 | 1.3 | 8.0 | 0.7 |
| COVID Lung Tissue [2] | 27 | 4,308 | 0.4 | 1.0 | 1.9 | 9.0 | 0.3 |
| Mouse Brain Tissue [3] | 59 | 19,817 | 341.8 | 9.3 | 151.7 | 58.5 | 1.0 |
| Colorectal Tumor [4] | 99 | 3,629 | 1.6 | 5.6 | 6.5 | 45.3 | 0.5 |
| COVID PBMC [5] | 143 | 4,527 | 1.8 | 10.0 | 11.6 | 60.0 | 0.4 |
| Lung Fibrosis [6] | 144 | 4,965 | 3.0 | 14.0 | 13.5 | 79.6 | 0.6 |
| Lupus PBMC [7] | 336 | 3,761 | 57.1 | 16.9 | 142.3 | 147.7 | 0.4 |

Table S1: **Timing (in minutes) of GloScope and Other Data Processing Steps** Each row provides the the runtime in minutes of GloScope on the given dataset, as well as the common scRNA-Seq processing steps of computing PCA, cell clustering, and UMAP. We provide both the total runtime of GloScope, across all samples, as well as the average runtime when considered per sample. The primary computational overhead of GloScope is fitting a Gaussian mixture model to each sample, and the total runtime is approximately linear in the number of samples. Because of their size, the processing of Lupus PBMC and Mouse Brain Tissue use computational tools for *SingleCellExperiment* objects in R/Bioconductor for the processing steps, whereas all other datasets utilize tools for *Seurat* objects (also in R). These experiments were run on machines with dual 32-core AMD EPYC 7543 processors and 512GB of RAM.

### Supplementary Figures

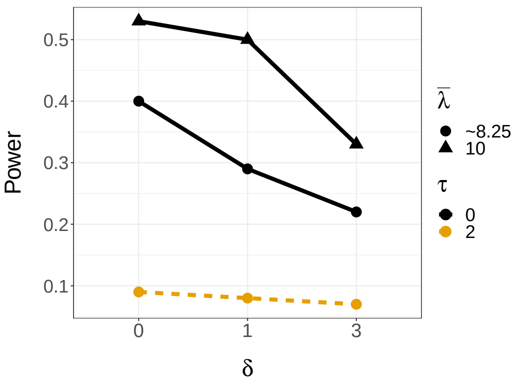


(A) Library size ($\lambda$)

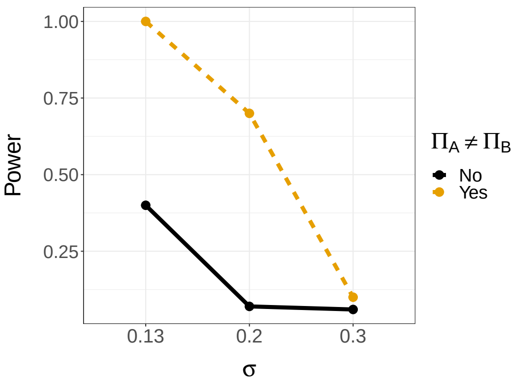


(B) Gene-level variability ($\beta$)

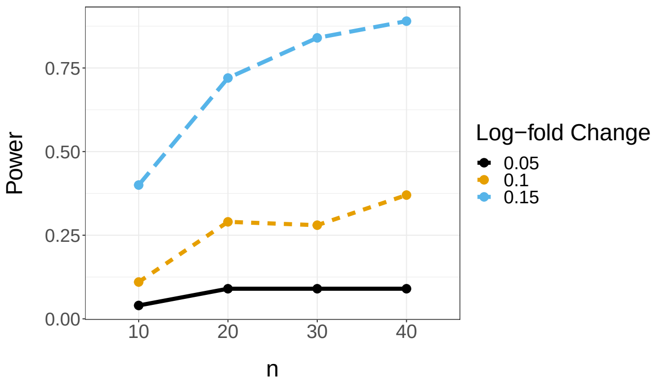


(C) Sample size ($n$ in each group)

Fig. S1: **Effect of changing various sources of sample variability on the power to detect group differences.** (A) Power to detect log-fold change differences in the presence of variation in the average library sizes between samples ($\lambda$) and individual cells within a sample ($\tau$); (B) Power to detect log-fold change differences in the presence of variation in the baseline expression levels between samples ($\sigma$); (A) and (B) have log-fold changes on average of 0.15 in 10% of DE genes. (C) Power to detect log-fold change differences in the presence of variation in the sample size within a single groups ($n$). Power of ANOSIM calculated based GloScope representation using GMM density estimation and reduced dimensionality representation via PCA with 10 dimensions.

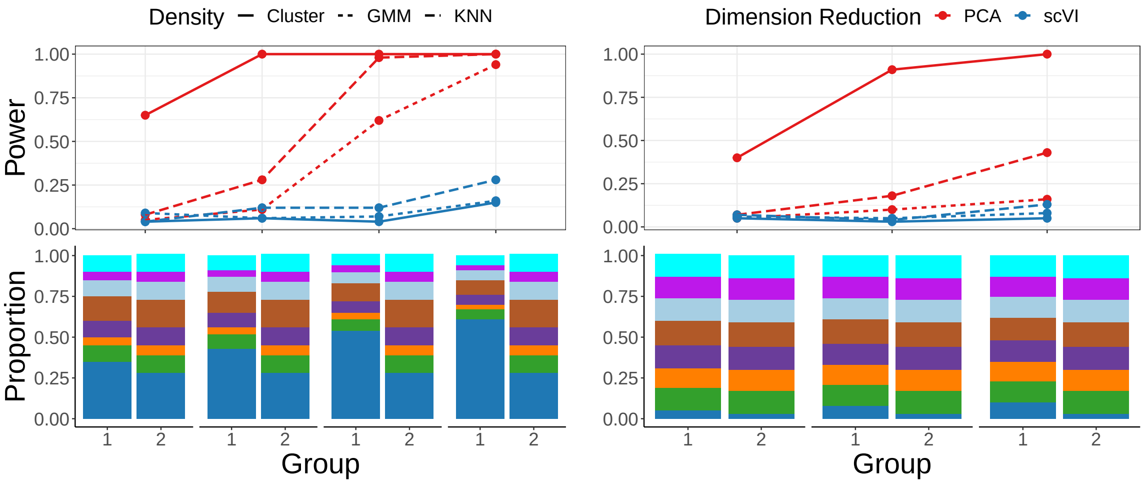


Fig. S2: **Change in cell-type composition (no DE genes).** Major changes were in the two groups' largest cluster (left) or smallest cluster (right). The cell-type composition is visualized in the lower panels. Each group consists of n=10 samples with m = 5000 cells per sample (the sample level variability parameter $\sigma$ is fixed at 0.13, and the sequencing depth $\lambda = 8.25$, see Methods for details on these parameters). Power calculated based on cluster proportion vector, GMM or kNN density estimation, and reduced dimensionality representation via PCA or scVI with 10 dimensions.

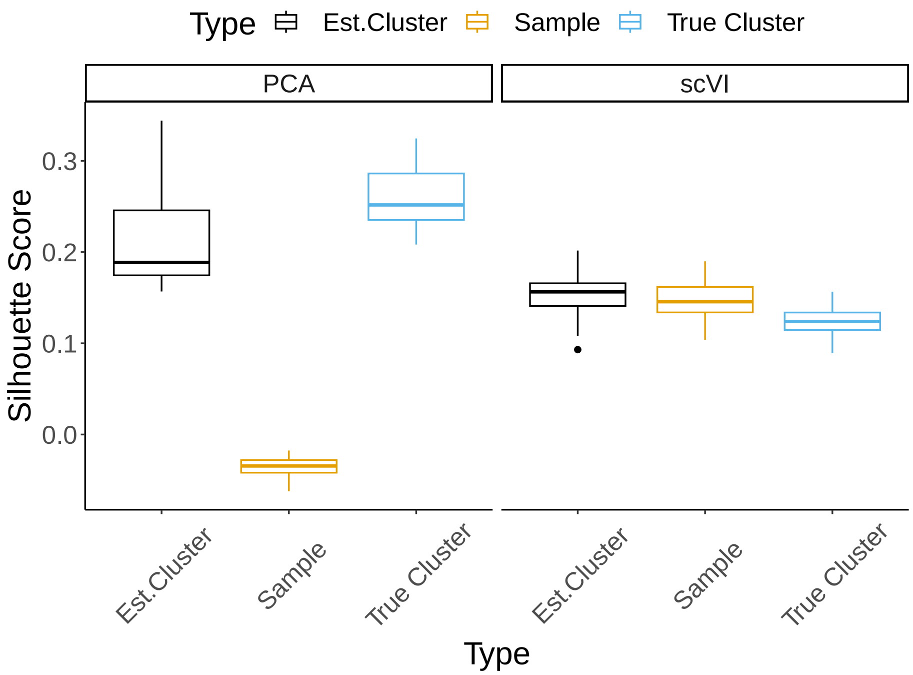


Fig. S3: **Evaluation of PCA and scVI discrimination of samples and group variability.** Individual cells were simulated from 10 sample with sample-level variability ($\sigma = 0.13$) and reduced to 10 dimensions, either with PCA or scVI. For each simulation, the silhouette score of the reduced dimensionality reduction was calculated at the individual cell-level to assess the similarity of cells within the same sample, compared to the similarity of cells within the same subtype. Larger values indicate larger separation between either samples or subtypes. PCA shows small variation between samples compared to the variation between subtypes, while Each boxplot consists of the silhouette scores for assessing the goodness of clustering different factors for dimension reduction embeddings obtained from either PCA or ScVI. 100 simulations were made to estimate the distance matrics.

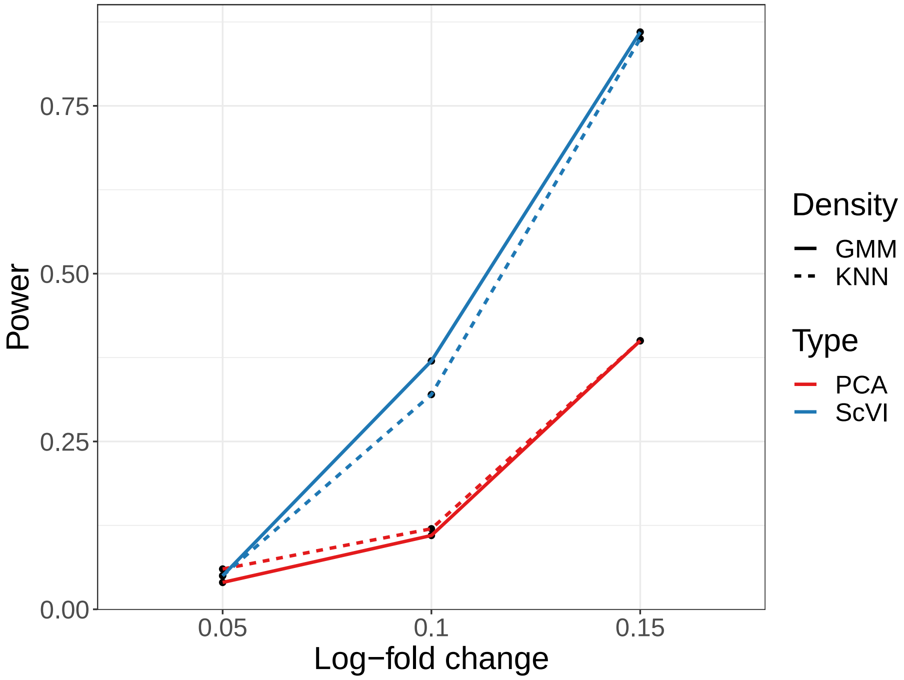


Fig. S4: **Evaluation of the different choices by calculating the power of detecting gene expressios.** 100 simulations were made to estimate the distance matrics. Power of ANOSIM calculated based GloScope representation using kNN or GMM density estimation and reduced dimensionality representation via scVI or PCA with 10 dimensions. ScVI shows much stronger power of between group difference detection compared to PCA, while there is not much distinction observed when compare GMM vs kNN.

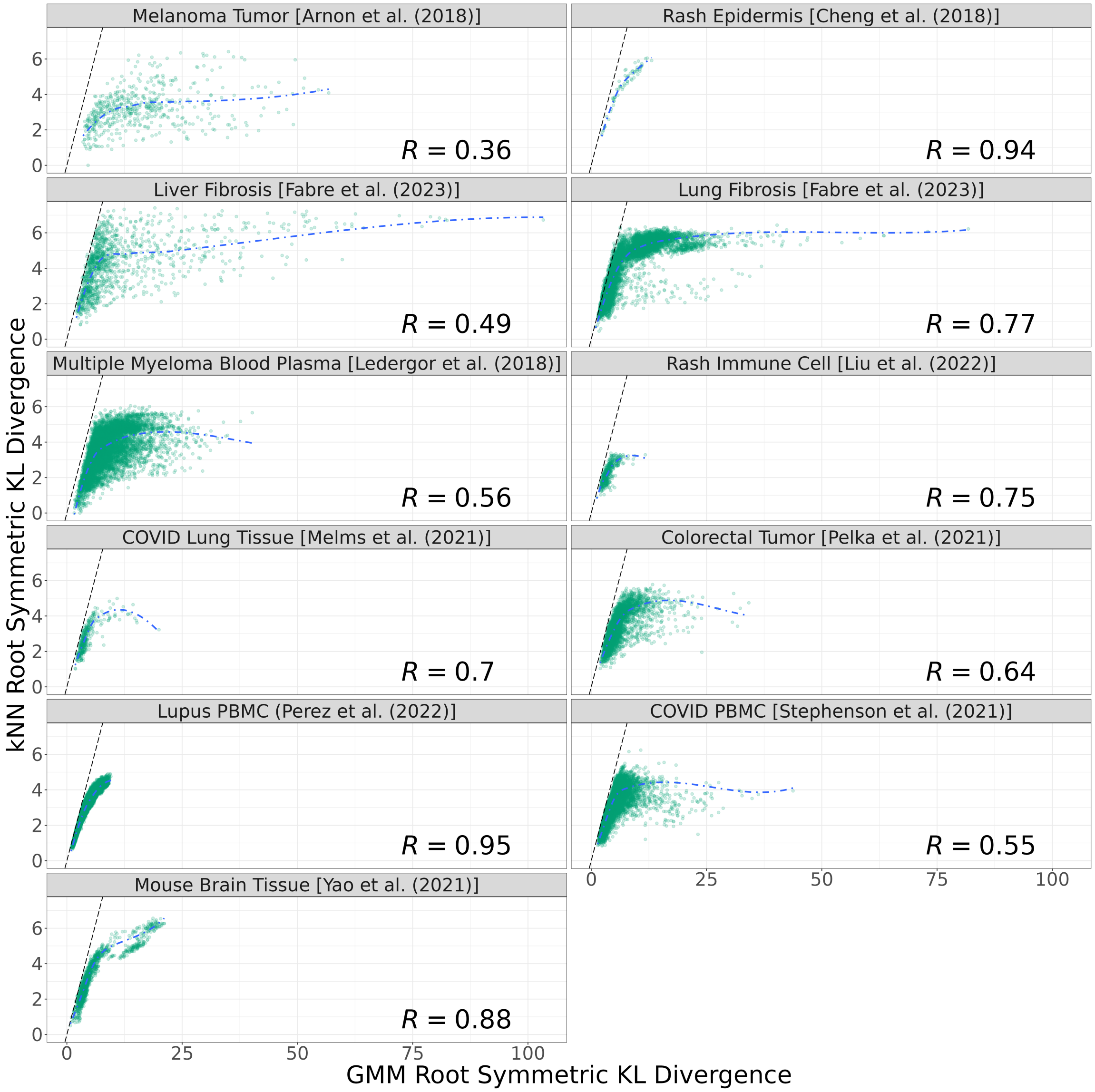


Fig. S5: **Sample-pair root divergences compared between GMM and kNN density estimators after PCA dimensionality reduction.** Each subplot summarizes the GloScope divergences between samples in one of the 11 datasets considered in this paper. Each point in the scatter corresponds a pair of samples. The x-axis coordinate is their symmetric KL divergence estimated from GMM densities, and the y-axis coordinate is that divergence estimated from kNN densities. A LOESS curve is fit to model the mean trend across all smaple pairs, and Pearson correlation coefficients are indicated. The identity line is also marked for reference.

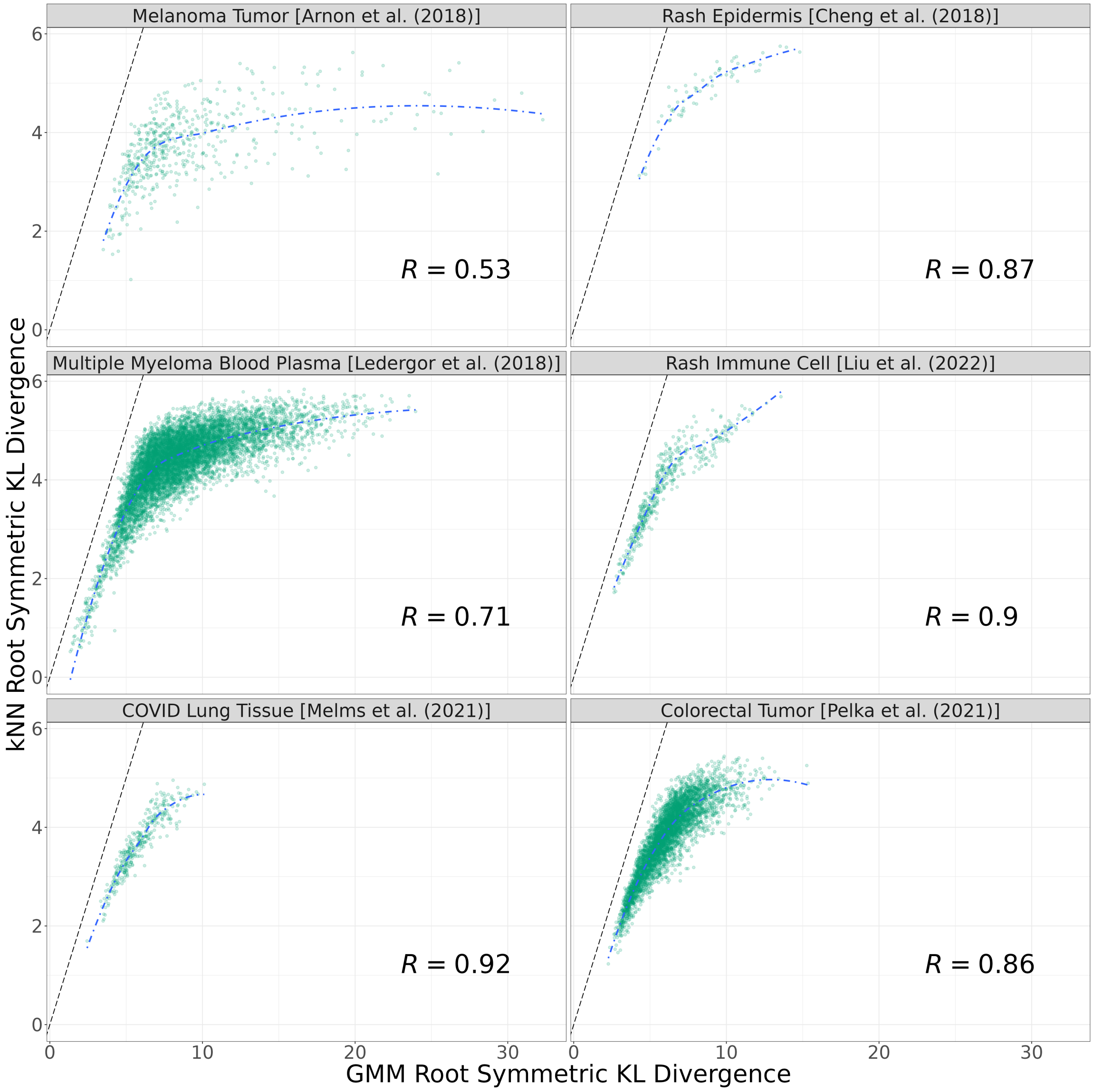


Fig. S6: **Sample-pair root divergences compared between GMM and kNN density estimators after scVI dimensionality reduction.** Due to computational limitations, scVI was only applied to 6 of the datasets we consider in this paper. See the caption of Figure S5 for information about the shared features of these scatter plots.

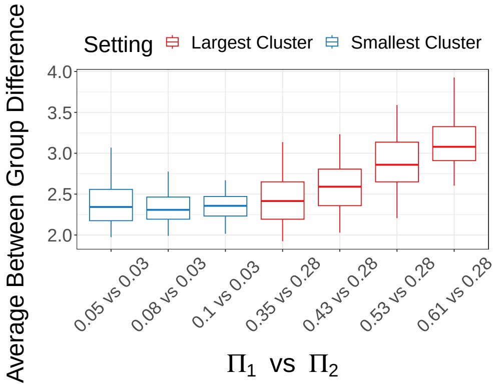


(A) GMM, PCA

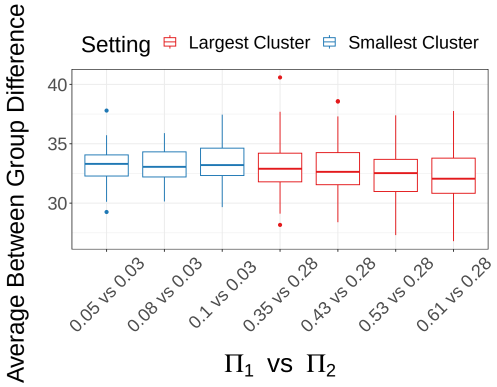


(B) GMM, scVI

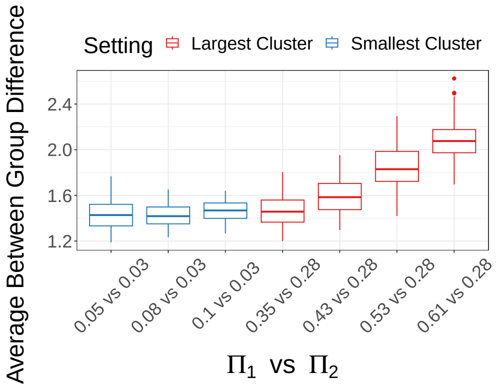


(C) kNN, PCA

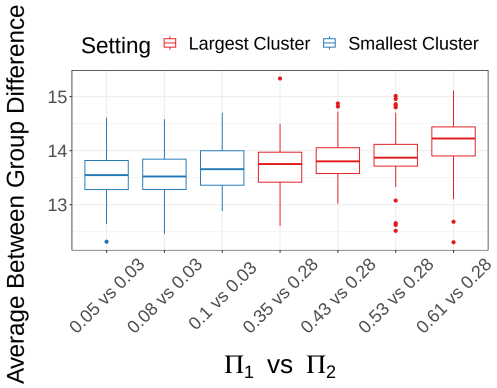


(D) kNN, scVI

Fig. S7: **Boxplot demonstration of global cell type composition changes detection by GloScope.** The major changes were in the two groups' largest cluster or smallest cluster (the actual values of the proportion changes in the largest or smallest group, $\Pi_1$ vs $\Pi_2$, are labeled in the legends). Each box is drawn from 100 simulation's average between group distance, calculated using 10 dim embeddings.

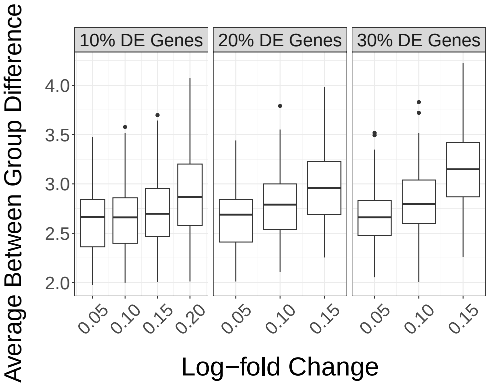


(A) GMM, PCA

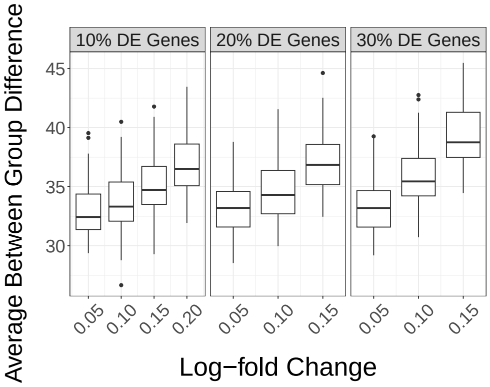


(B) GMM, scVI

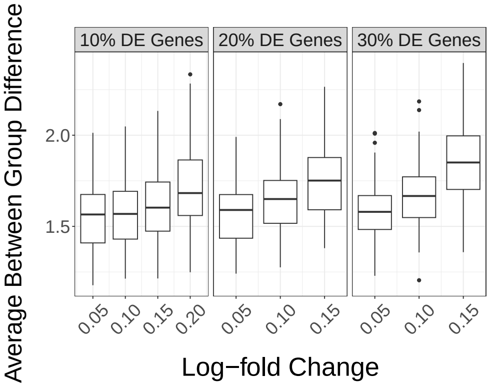


(C) kNN, PCA

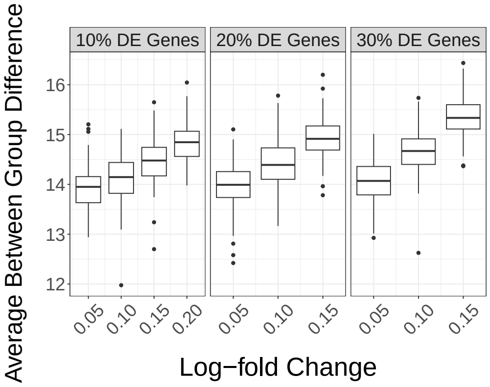


(D) kNN, scVI

Fig. S8: **Boxplot demonstration of gene expression changes detection by GloScope.** Each box is drawn from 100 simulation's average between group differences, calculated using either GMM or kNN density estimation with either 10 dimensional PCA or scVI 10 embeddings. Upward trend of distance was observed in each combination when log-fold change and percentage of DE genes increase.

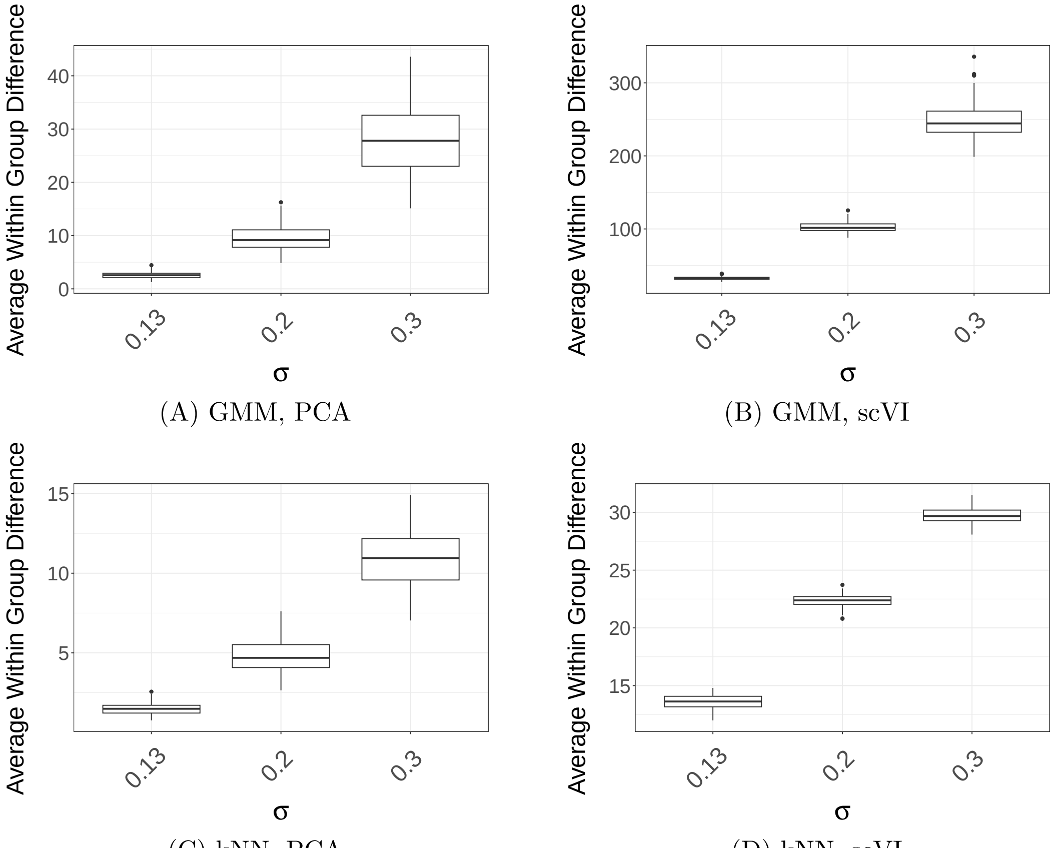


(A) GMM, PCA

(B) GMM, scVI

(C) kNN, PCA

(D) kNN, scVI

Fig. S9: **Boxplot demonstration of detecting increased sample level variation in the gene expression differences by GloScope.** Each box is drawn from 100 simulations' average divergences among sample within a single phenotype group distance using either GMM or kNN density estimation with either 10 dimensional PCA or scVI 10 embeddings. 10 dimensions. Larger variation of average within group distance could be easily detected in most combination when sample level gene expression variation $\sigma$ increases.

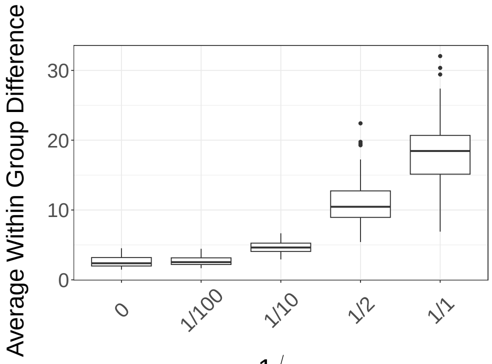


(A) GMM, PCA

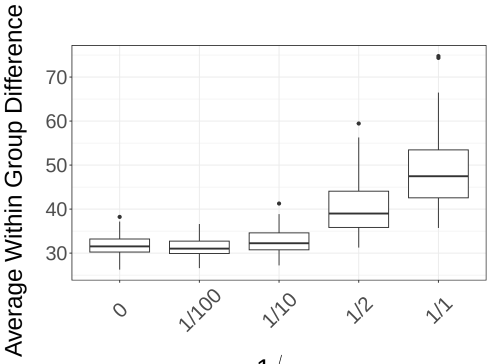


(B) GMM, scVI

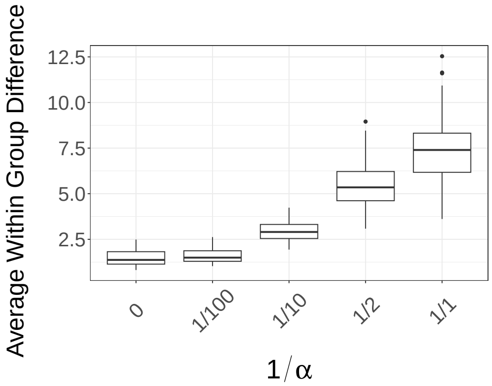


(C) kNN, PCA

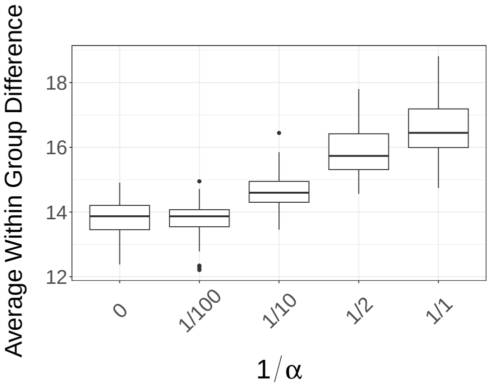


(D) kNN, scVI

Fig. S10: **Boxplot demonstration of detecting increased cluster proportion variation $\alpha$ by GloScope.** Each box is drawn from 100 simulations' average divergence among samples within a single phenotype group, calculated using eitherr GMM or kNN density estimation with either 10 dimensional PCA or scVI 10 embeddings. Larger variation in the average within group distances can be easily observed When sample level cluster proportion variation $1/\alpha$ gets larger.

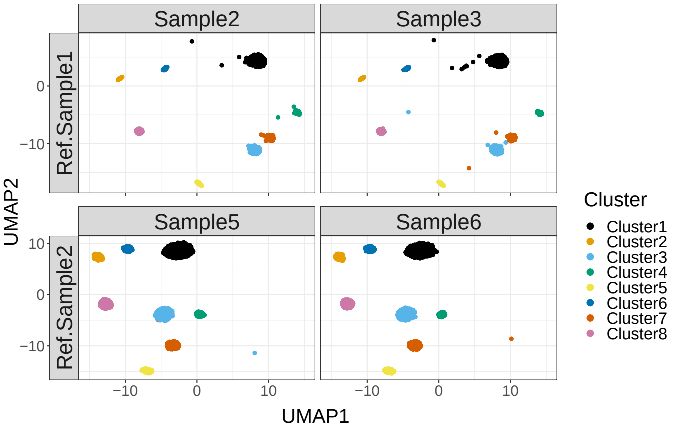


(A) UMAP representation for 1 simulation run with original muscat pipeline

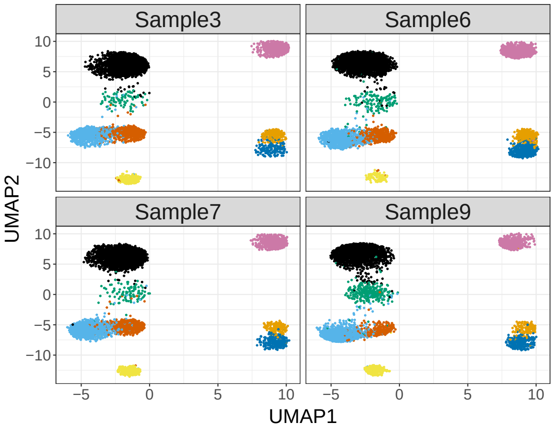


(B) UMAP representation for 1 simulation run with modified simulation pipeline

Fig. S11: **UMAP plot demonstration of original muscat simulation pipeline versus modified simulation pipeline.** (A) shows the umap representation of simulated data from original muscat pipeline, where strong sample batch was observed: samples from first row was simulated from the same reference sample and sample from the second row was simulated from the same reference sample. (B) shows that after modifying $\beta_k$, some clusters were brought closer to or mixed with each other, and remove the strong sample batch due to the recycled parameters. Such modification allows the simulated data to have more reasonable and similar behavior to the real scRNA-Seq data than the data simulated using muscat pipeline.

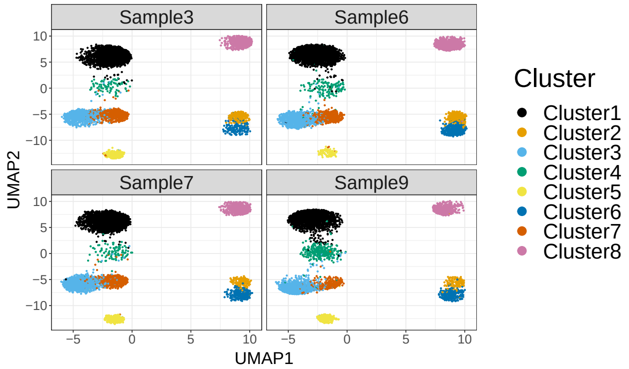


(A) UMAP representation for 1 simulation run with $\sigma = 0.13$

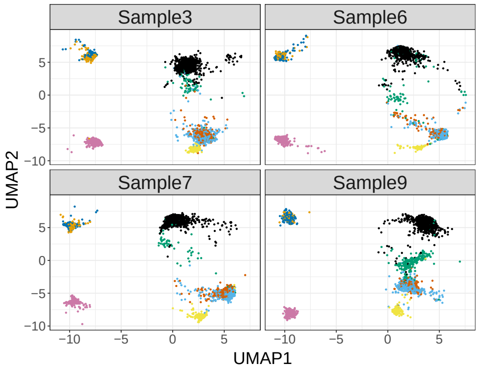


(B) UMAP representation for 1 simulation run with $\sigma = 0.3$

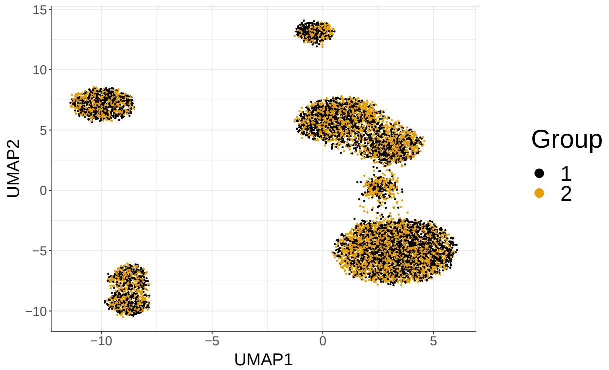


(C) UMAP representation for 1 simulation for lfc = 0.05

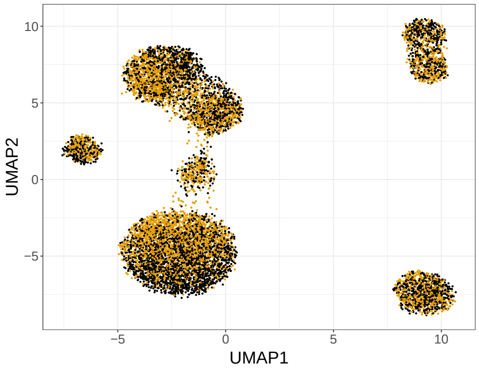


(D) UMAP representation for 1 simulation for lfc = 0.15

Fig. S12: **UMAP plot demenstration of different parameter effects, including gene expression changes and sample level variation.** Each plot is drawn from 1 particular simulation realization. (B) shows that increasing $\sigma$, the gene expression level variation, leads to more varied expression among samples compared to (A). (D) shows the increased log-fold change effect compared to (C).

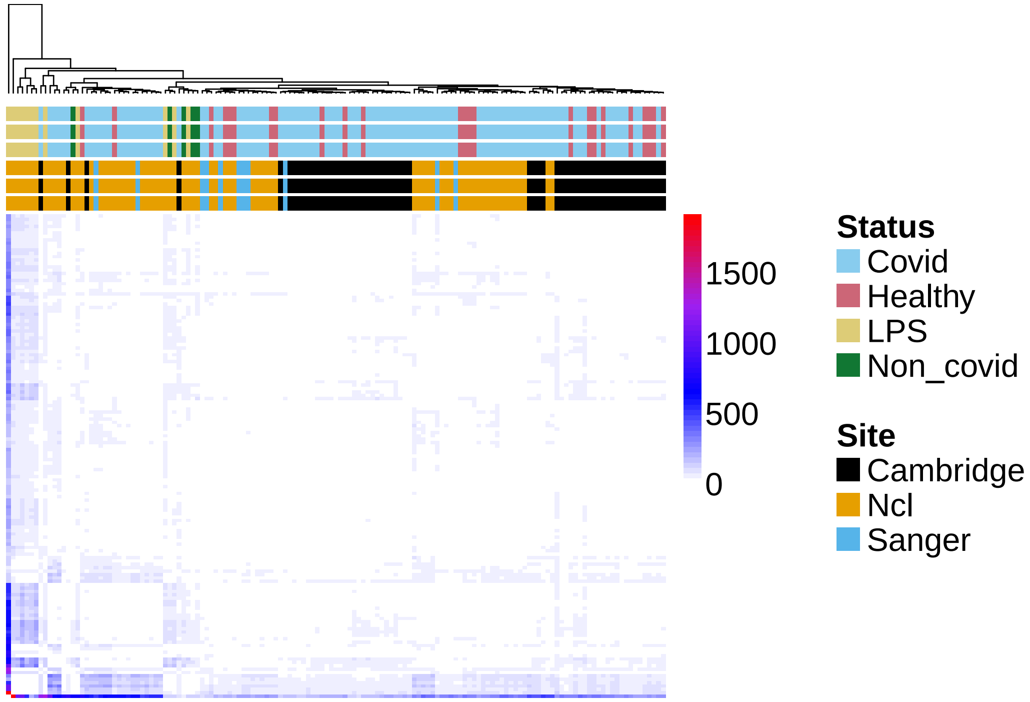


Fig. S13: **Full distance matrix of the GloScope estimates of divergence of the original COVID PBMC single-cell data.** Shown is a heatmap of the pairwise divergences calculated by GloScope for the 143 samples of [5]. Samples are diagnosed from COVID, healthy control, volunteer administered with LPS stimulus and the patients suffering from other non-COVID respiratory disease, and their phenotype is noted at the top of the heatmap. Estimated distances used the GMM estimate, and latent variable estimated with PCA with 10 dimensions.

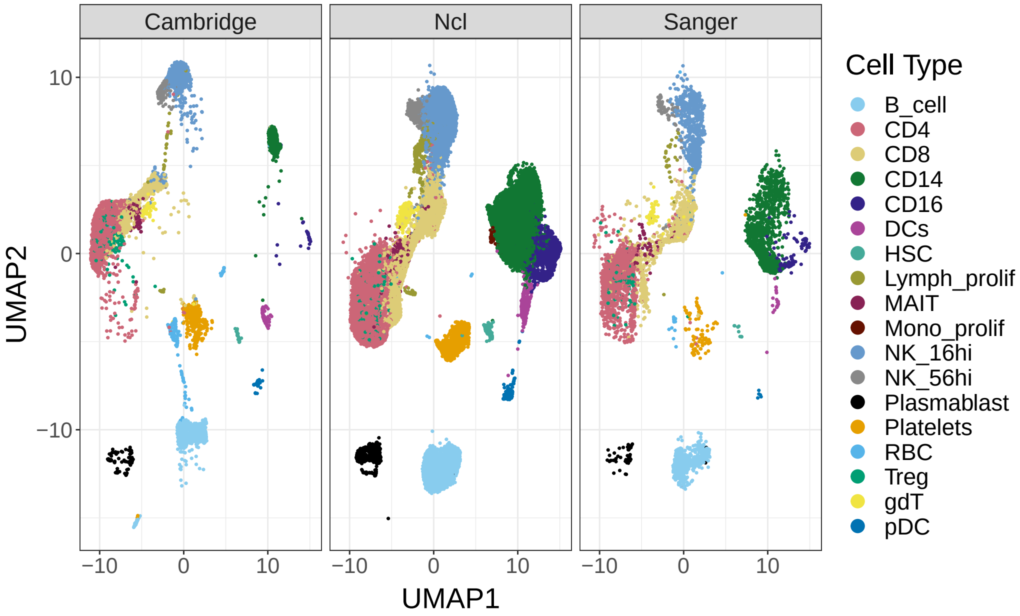


Fig. S14: **UMAP plot of the original COVID PBMC single-cell data [5].** Each panel of the plot contains all cells from the same sequencing site, regardless of sample or disease status. The cells are color-coded in each panel by the cell-type of the cell, as identified by Stephenson et al. [5] following batch correction with Harmony. The UMAP embedding was calculated from the first 30 PCA dimensions using all cells, and Only a random subset of 50,000 cells were selected for plotting in the figure. This visualization shows the clear differences due to sequencing site in the cells which were identified to be in the same subtype, such as B-cells, CD4 cells and NK_56 high (CD56 bright NK cells).

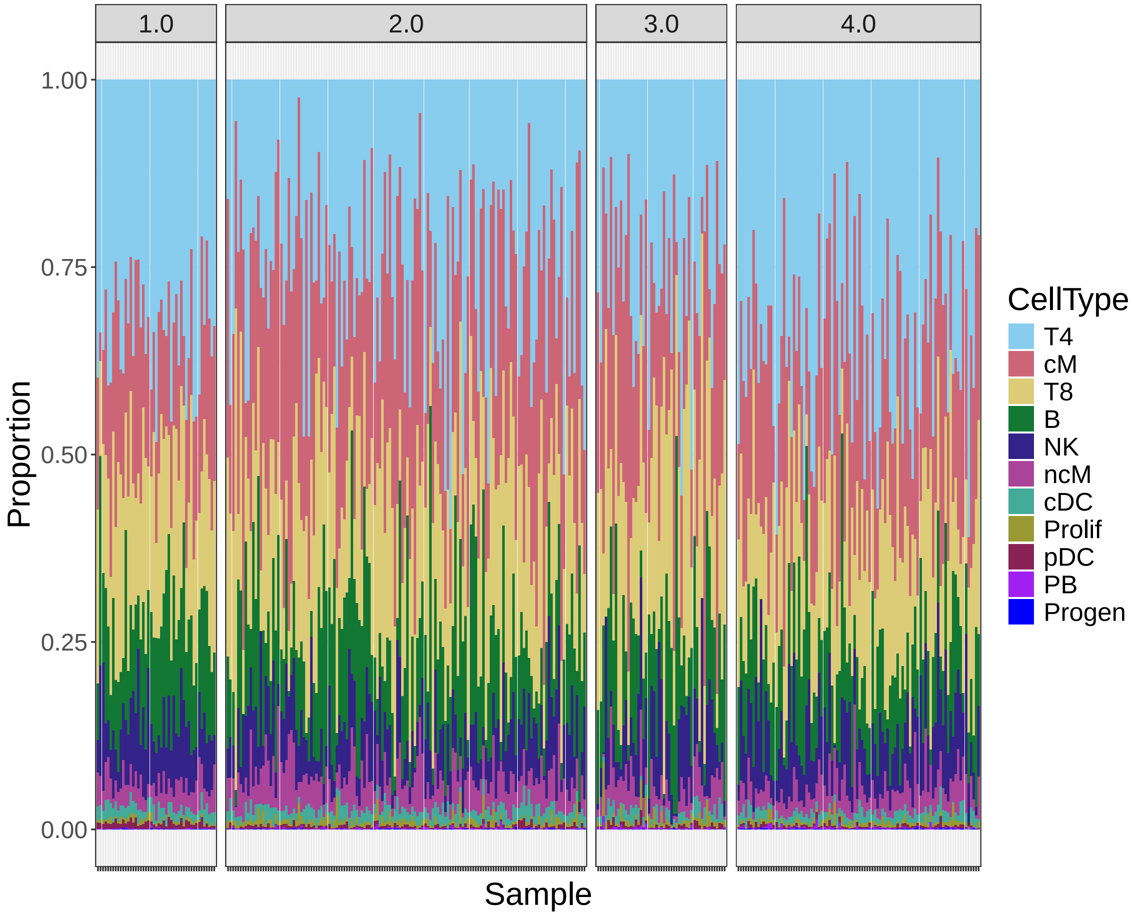


Fig. S15: **Bar plot visualization of cell type proportion per samples in the original batches of the Lupus PBMC study [7].** For plot details and color annotation, see Fig. S16. Panels are separated by original batch annotated by the Perez et al. [7], without further separation of batch 4.0 into subgroups identified by GloScope.

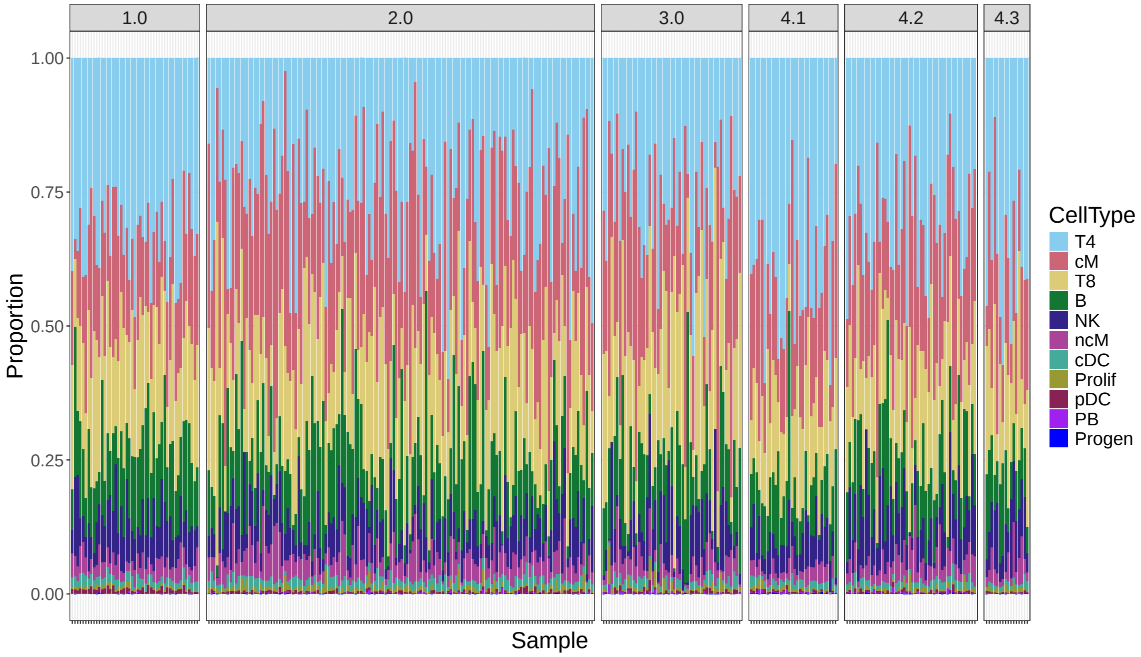


Fig. S16: **Barplot visualization of cell-type proportion differences in subgroups identified by GloScope for Lupus PBMC study [7].** Each column/bar represents a sample. The bars are broken into different color-coded segments, with a segment for each cell-type and the size of the segment proportion to the proportion of cells in the data identified with the cell-type. The annotation of individual cells into cell-types are based on the annotation provided by Perez et al. [7] using canonical marker genes. Samples are separated in different panels based on their processing batches provided in Perez et al. [7], with the *de novo* subgroups found by GloScope in the fourth processing batch shown separately. For the subgroups of the fourth processing batch, we see samples in batch 4.1 has relatively larger proportion of CD4 T cells than batch 4.2 and 4.3.

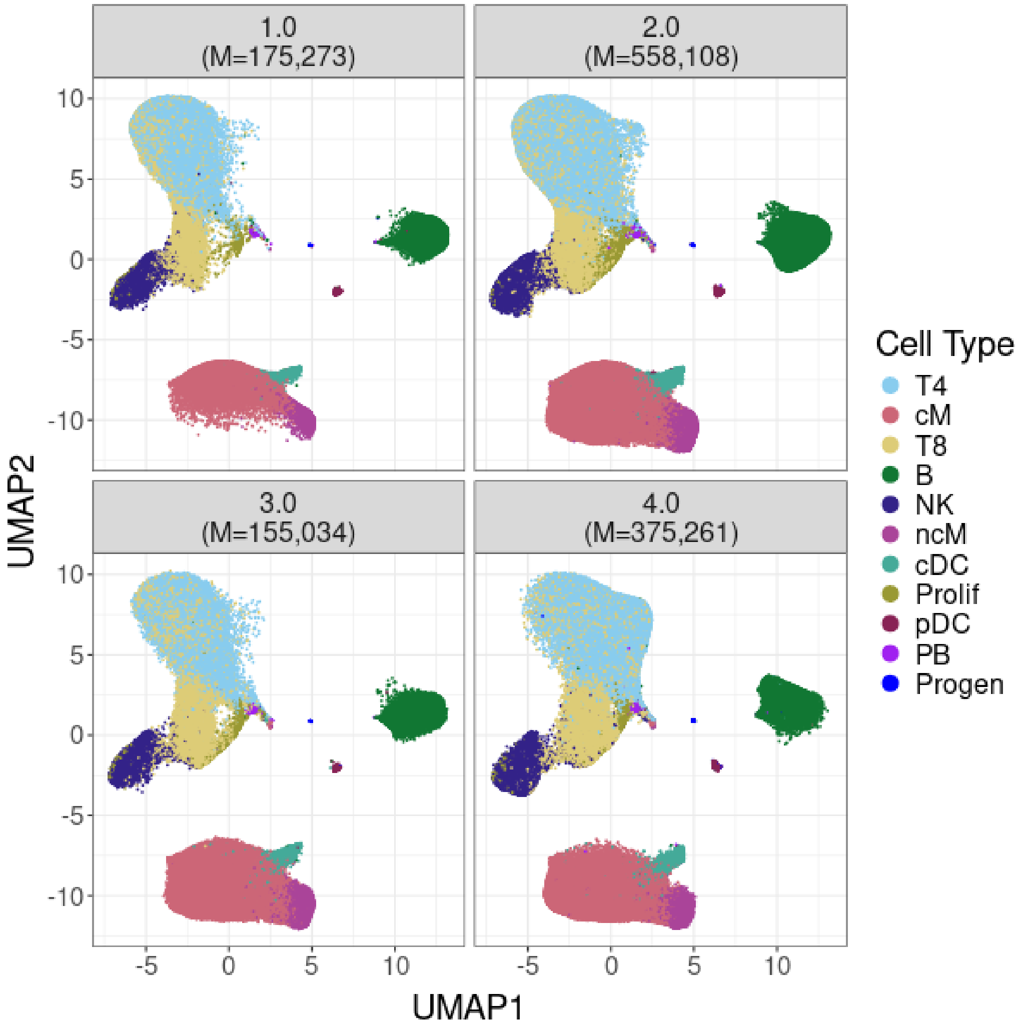


Fig. S17: **UMAP visualization of the gene expression per batch of data from Perez et al. [7].** Cells are separated in different panels for batches provided by Perez et al. [7]. The number of cells ($M$) plotted in each panel are indicated in the panel title. The cells are color-coded in each panel by the cell-type identified by Perez et al. [7] using canonical marker genes. The first 10 PCs were used to calculate the UMAP representation across all samples.

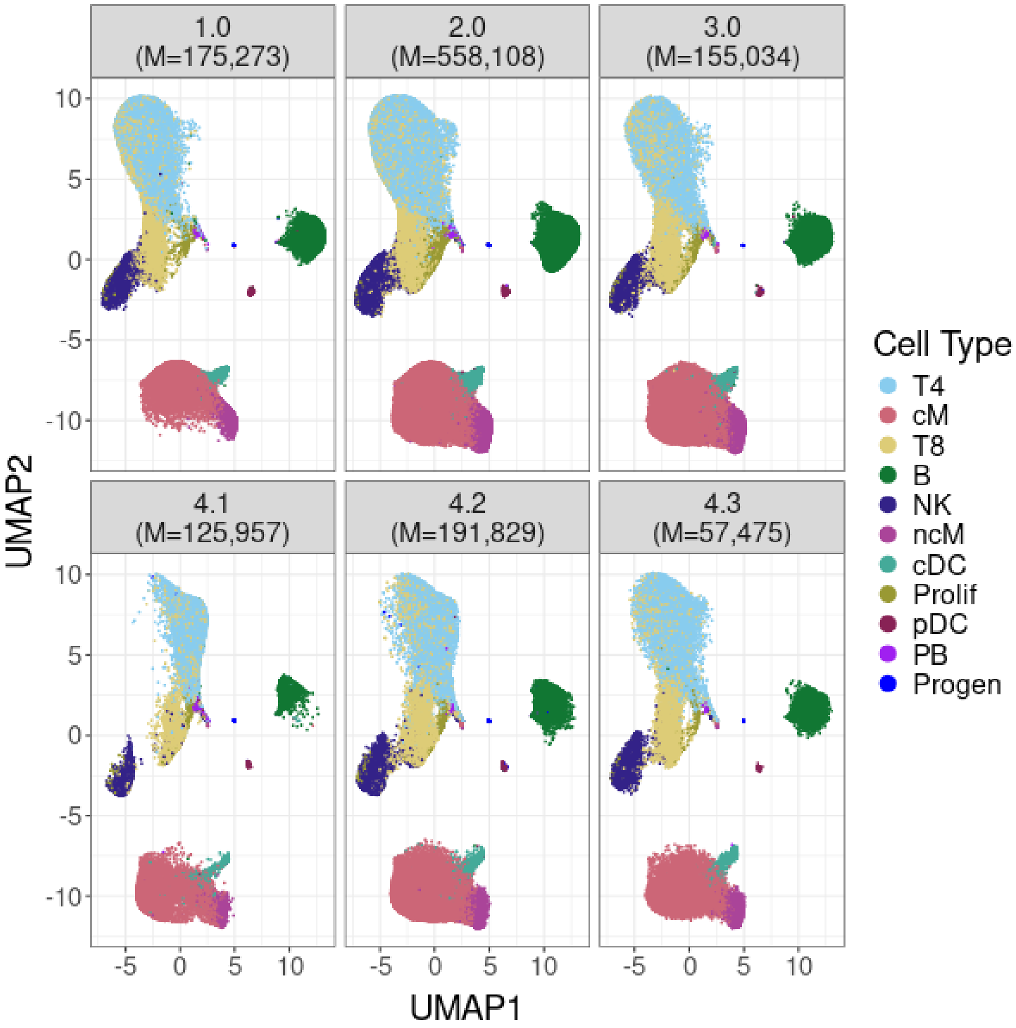


Fig. S18: **UMAP visualization of the potential subgroups of batch 4 from Perez et al. [7].** For description of UMAP calculations and color annotations, see Fig. S17. The upper panel is the first 3 original processing batches provided by Perez et al. [7], as shown in Fig. S17. The lower panel further separates the fourth batch into the subgroups identified by GloScope.

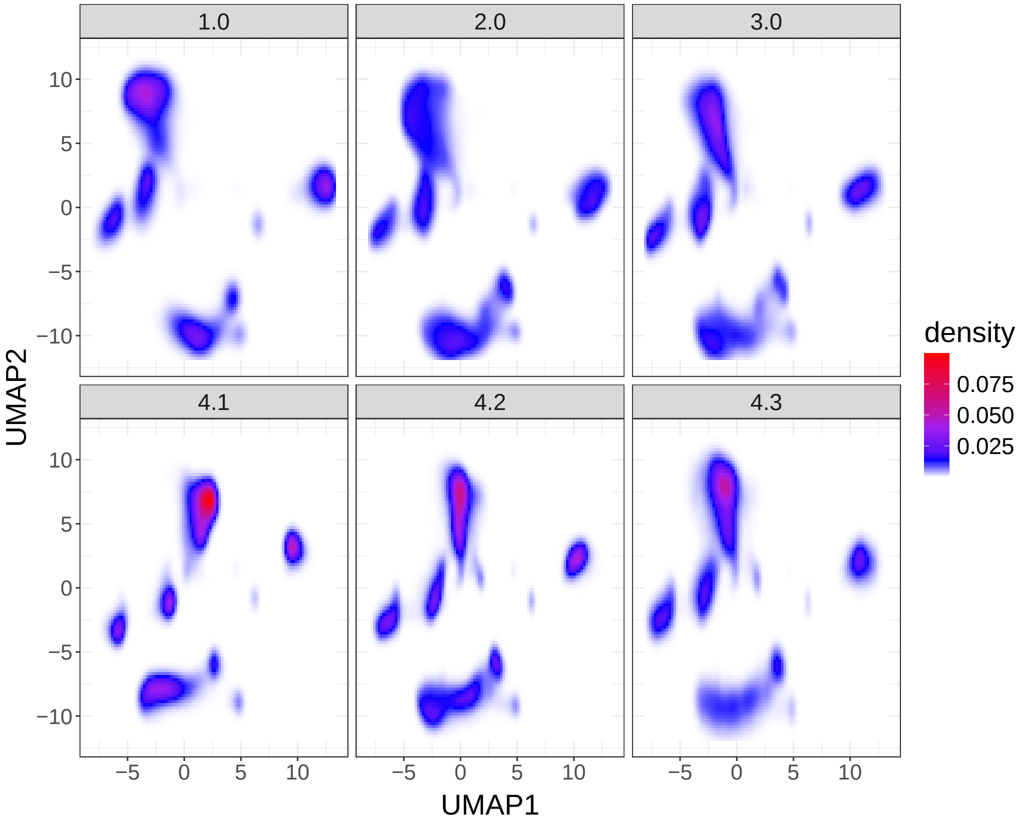


Fig. S19: **Visualization of the cell density representation of the UMAP in Fig. S17.** For UMAP calculation and panel explanation, see Fig. S18. Instead of plotting the individual cell UMAP coordinates, the UMAP embeddings were used to create a 2-dimensional density estimate, and plotted in this figure based on a color gradient shown in the legend. Plotting the density values allows us to see shifts in distribution between batches 1-3 that are not easily apparent due to overplotting in Fig. S18. Even with a density plot, it can be difficult to align the densities across multiple panels, further demonstrating the difficulty in drawing conclusions about shifts from UMAP-based visualizations.

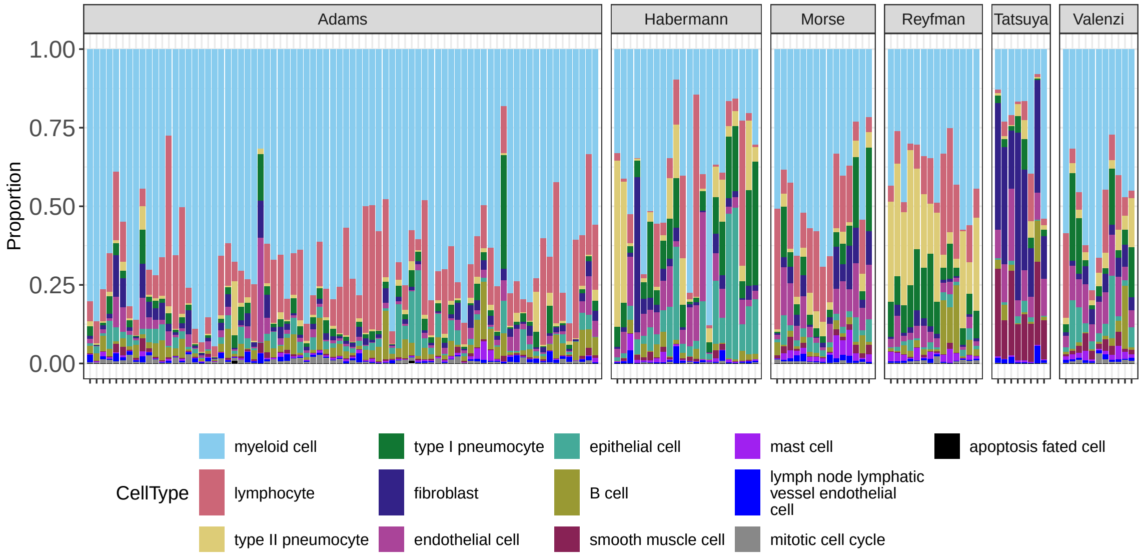


Fig. S20: **Barplot visualization of celltype proportion per sample of lung study in Fabre et al. [6].** Each column represent a sample and grouped into different panels by the study where the samples were collected. Bars are color-coded by the cell types identified by Fabre et al. [6] following batch correction with Harmony. We are able to detect significant cell proportion differences (e.g myeloid cells) between Adams et al. [8] and other studies.

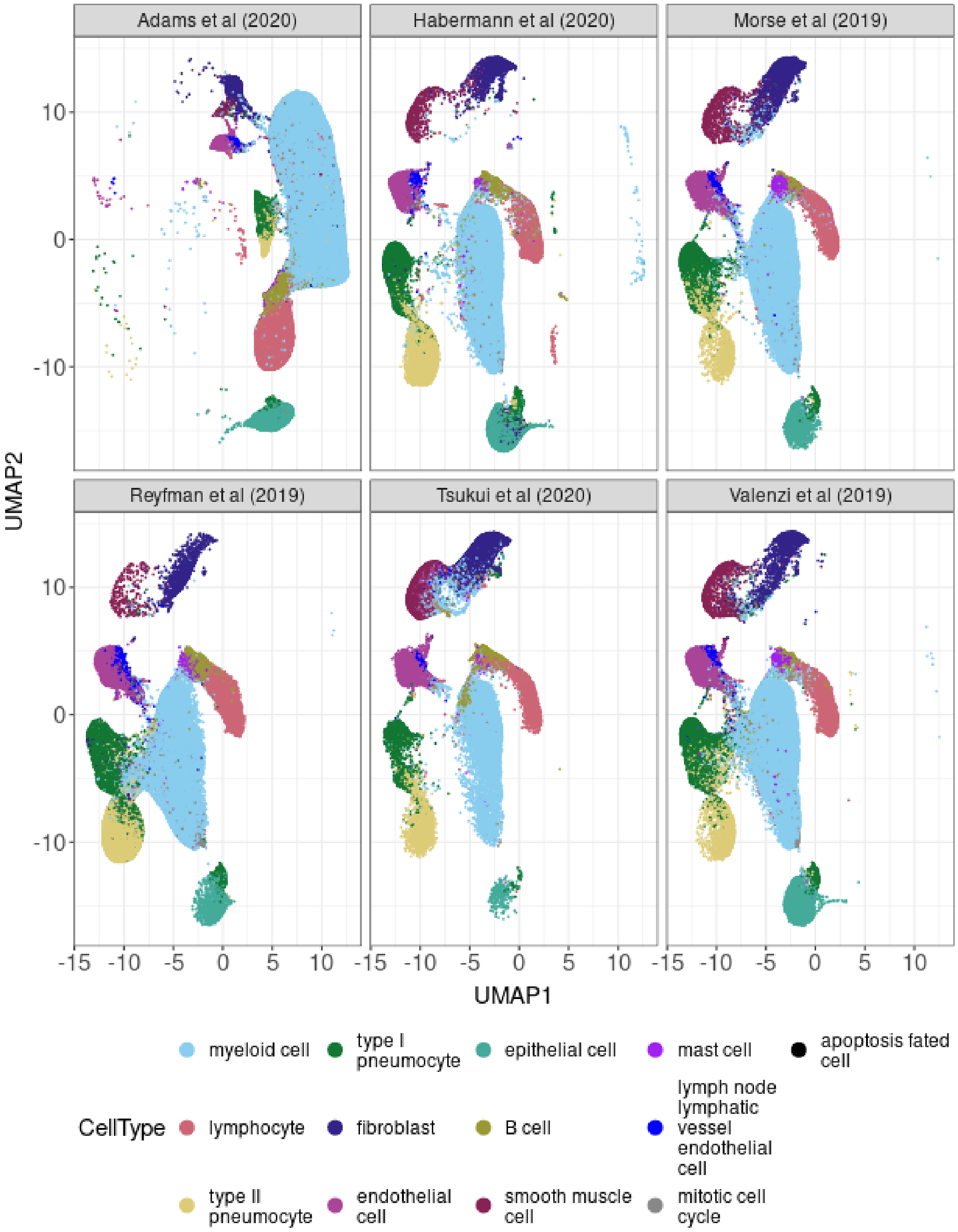


Fig. S21: **UMAP visualization of individual cells from lung study in Fabre et al. [6].** Each panel corresponds to cells in the six studies being integrated by Fabre et al. [6], with Adams et al. [8] showing widespread differences from the other studies. The cells are color-coded in each panel by the cell-type identified by Fabre et al. [6] following batch correction with Harmony. The first 10 PCs calculated on all the cells jointly are used for UMAP calculation.

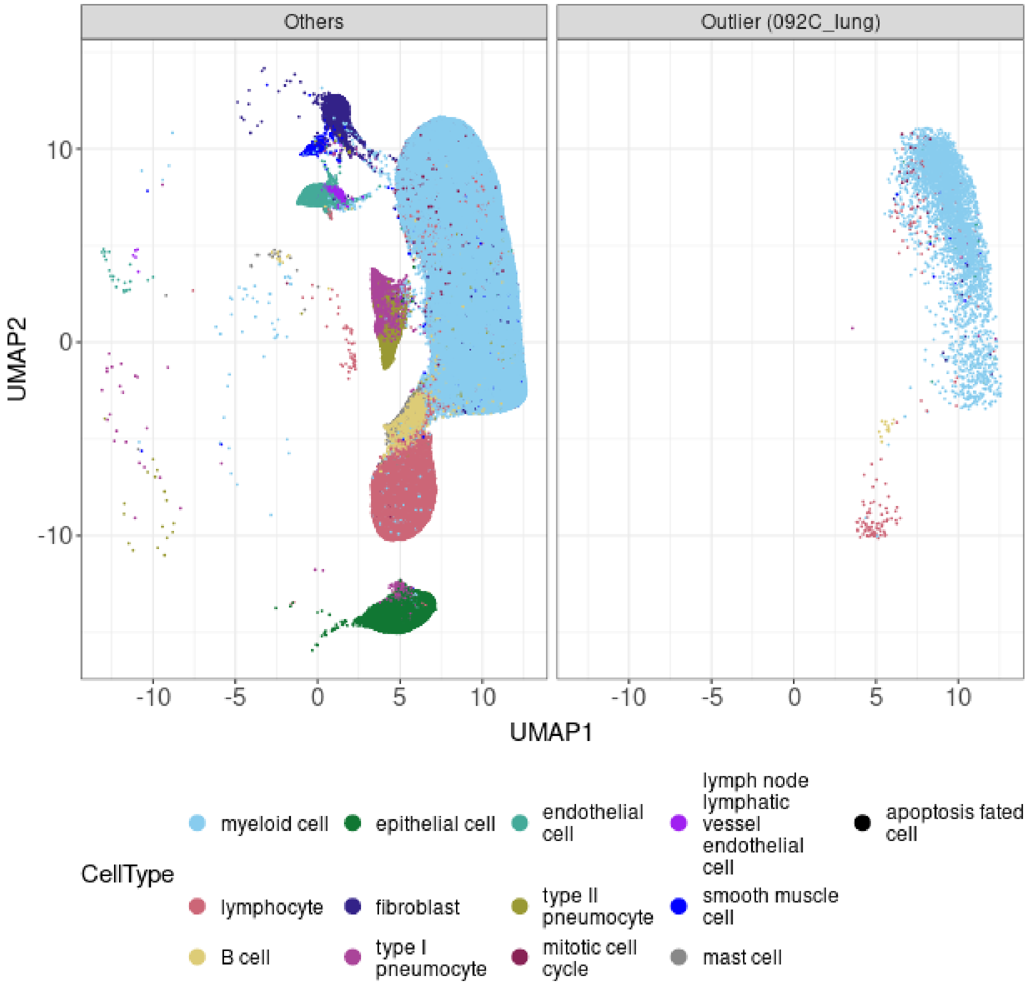


Fig. S22: **UMAP visualization of individual cells of outlier sample compared to other samples from Adams et al. [8].** For UMAP calculation and color annotation, see Fig. S21. Left panel is the cells from Adams et al. [8] where the samples are not considered as outliers, and right panel is the cells from the outlier sample (092C_lung). Most of the cell types are missing for the outlier samples

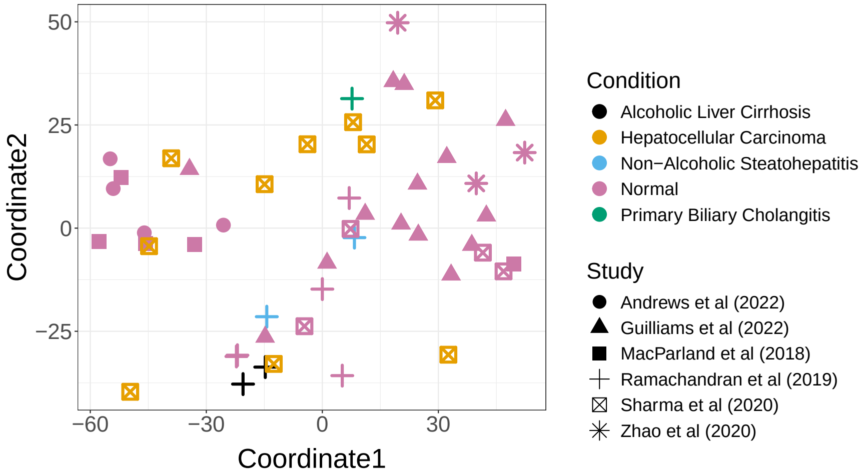


(A) MDS Plot

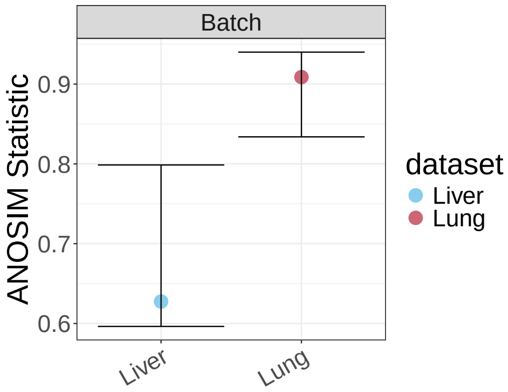


(B) ANOSIM Statistics

Fig. S23: **Visualization of sample from liver study in [6].** (A) A MDS plot of the divergences calculated by GloScope, with samples color-coded by their biological condition and with the shape of the point indicating the study of origin. The liver study shows less obvious study effects compared to lung study. (B) Comparison of the ANOSIM Statistic ($R$) based on GloScope divergences to quantify the separation between samples in different studies for both the liver and lung studies; larger values of $R$ indicate more separation between groups. Individual points show the ANOSIM statistic, with bootstrap confidence intervals indicated by whiskers.

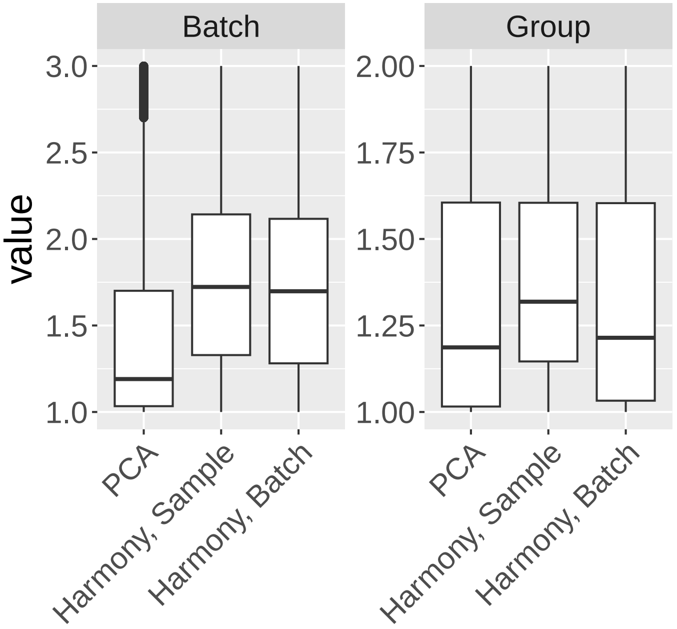


Fig. S24: **Boxplot of iLISI value for individual cells for data in Stephenson et al. [5].** The left panel showed the changes of iLISI value of each cell for batch quantification: the closer the values to 1, the more clear batch separation, indicating significatn batch effects; the closer the values to 3 (i.e. the number of batches), the better mixture among cells, indicating better batch correction. We saw that after applying Harmony on sample id and batch id, the iLISI values increased, suggesting the effectiveness of Harmony. The right panel showed the changes of iLISI value of each cells for separation of biological signal (COVID vs Healthy).

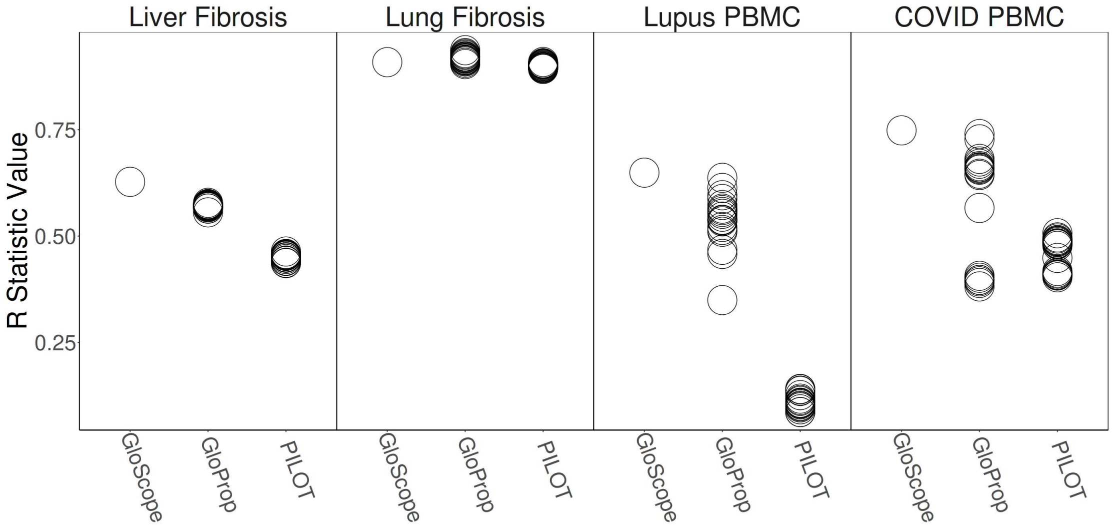


Fig. S25: **Detection of batch effect with GloScope, GloProp, and PILOT.** This figure compares how well sample-level represenations from GloScope, GloProp, and PILOT separate known batch effects in two fibrosis datasets [6], the lupus PBMC dataset [7], and the COVID PBMC dataset [5]. The batch separation statistic (y-axis) considered is the `ANOSIM` rank statistic ($R$) introduced by Somerfield et al. [9]. The replicated points for GloProp and PILOT denote 20 different random initializations used for the Leiden clustering algorithm with the resolution parameter $10^{-5}$. In all four datasets, GloScope is equivalent or improves upon the best performing example of GloProp and PILOT. Furthermore, the cluster composition-based methods exhibit considerable variability with the lupus PBMC and COVID PBMC data due to the random initalization.

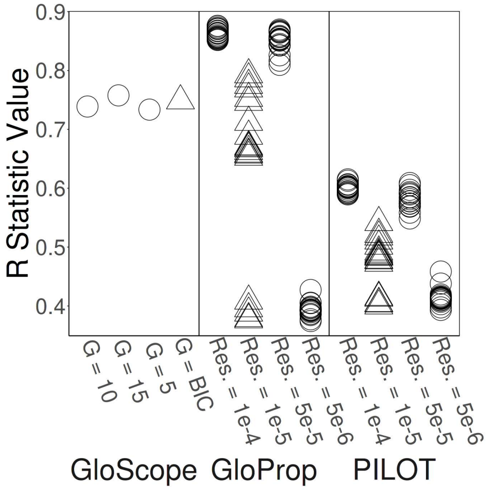


(A) COVID PBMC [5], $R$ Statistic

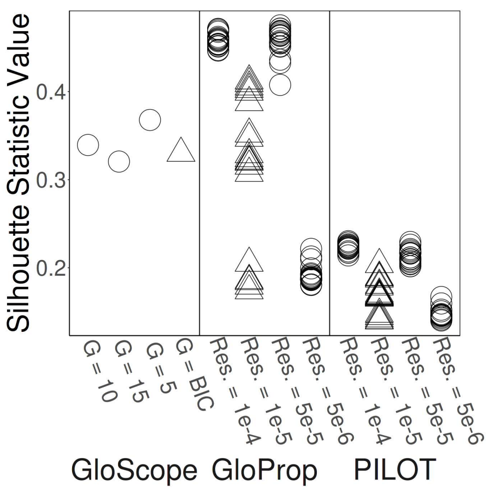


(B) COVID PBMC [5], Silhouette

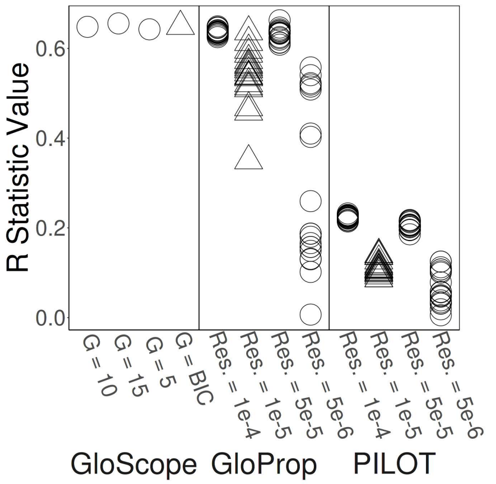


(C) Lupus PBMC [7], $R$ Statistic

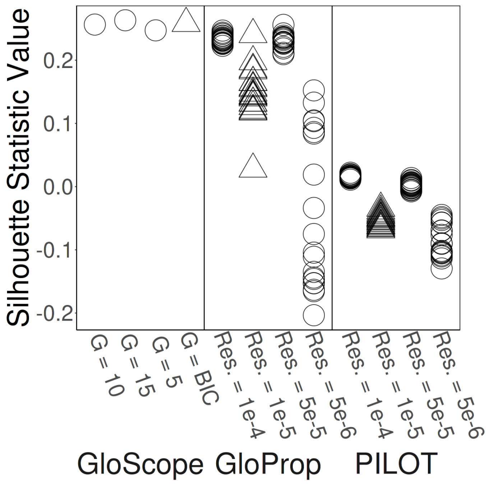


(D) Lupus PBMC [7], Silhouette

Fig. S26: **Demonstration of the variability due to parameter choices.** We demonstrate the effect of different parameter choices for GloScope, GloProp and PILOT. The y-axis shows the measured separation between the batches based on the (A),(C) `ANOSIM` $R$ Statistic, and (B),(D) based on the average silhouette width. (A) and (B) present the COVID PMBC data of [5]; (C) and (D) present the lupus PBMC data of [7]. For GloScope the parameter choices were the number of mixture components specified in the GMM density estimation step, including a grid search over 1 to 9 clusters to maximize Bayesian information criterion (BIC), the default in the `GloScope` package. For GloProp and PILOT, the parameter varied was the resolution (Res.) of the input Leiden clustering which assigns cluster labels to cells. Each column of points for GloProp and PILOT indicate a different resolution, and individual points within the same column represent the result of different random starts for the algorithm with a fixed resolution.

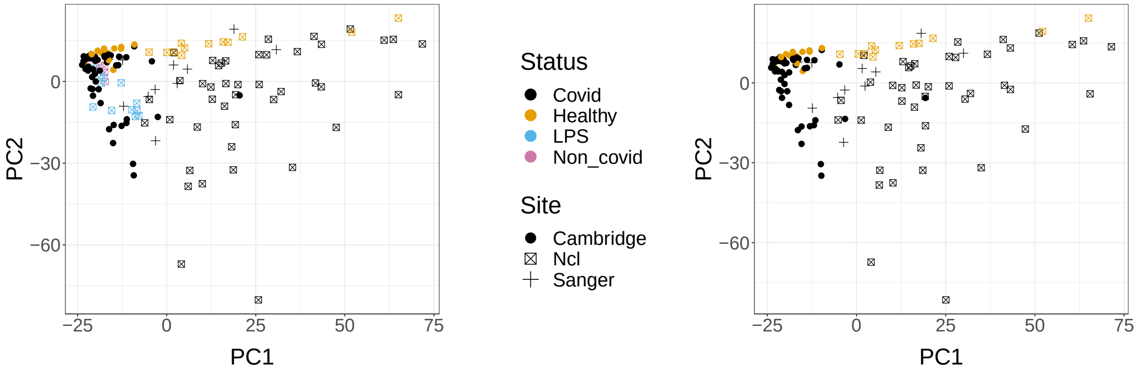


(A) Pseudobulk analysis on samples from COVID-19 study

(B) Pseudobulk analysis on COVID and Healthy samples from COVID-19 study

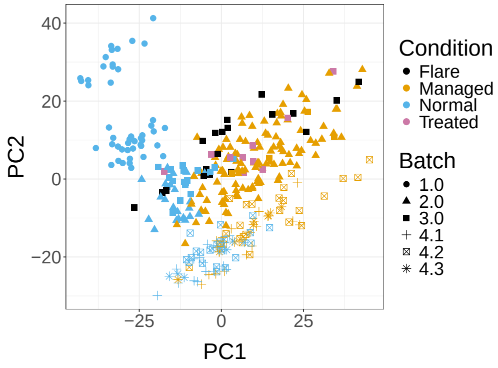


(C) Pseudobulk analysis of samples from lupus PBMC dataset

Fig. S27: **Visualization of the first 2 PC components of the pseudobulk.** (A) samples from COVID PBMC study of Stephenson et al. [5]. (B) Covid and Healthy samples from COVID PBMC study of Stephenson et al. [5]. Removing LPS and non-COVID samples yield similar results as in (A). (C) samples from lupus PBMC study of Perez et al. [7]. Note that the PCA coordinates are equivalent to performing the MDS on the matrix of pair-wise Euclidean distance between the samples.

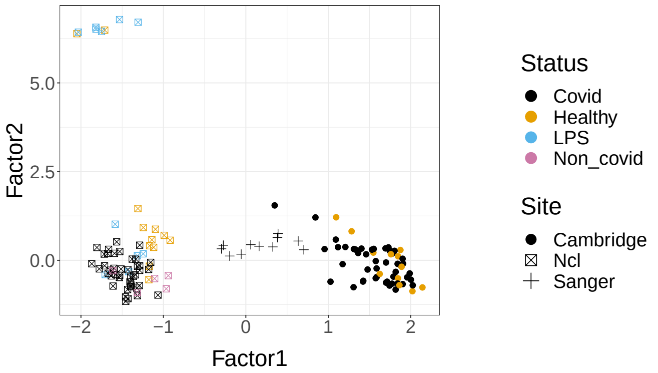


(A) MOFA on samples from COVID-19 study

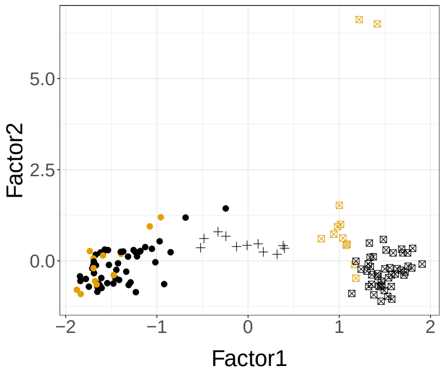


(B) MOFA analysis on COVID and Healthy samples from COVID-19 study

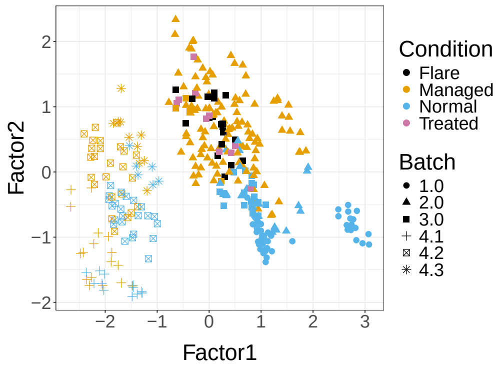


(C) MOFA analysis of samples from lupus PBMC dataset

Fig. S28: **Visualization of the first 2 factors of the MOFA results for data in Stephenson et al. [5] and Perez et al. [7].** (A) samples from COVID PBMC study of Stephenson et al. [5]. (B) Covid and Healthy samples from COVID PBMC study of Stephenson et al. [5]. Removing LPS and non COVID samples yield similar results as in (A). (C) samples from Lupus PBMC study of Perez et al. [7]. Each point is a sample, color-coded by their biological condition and with different shapes corresponding to their batch.

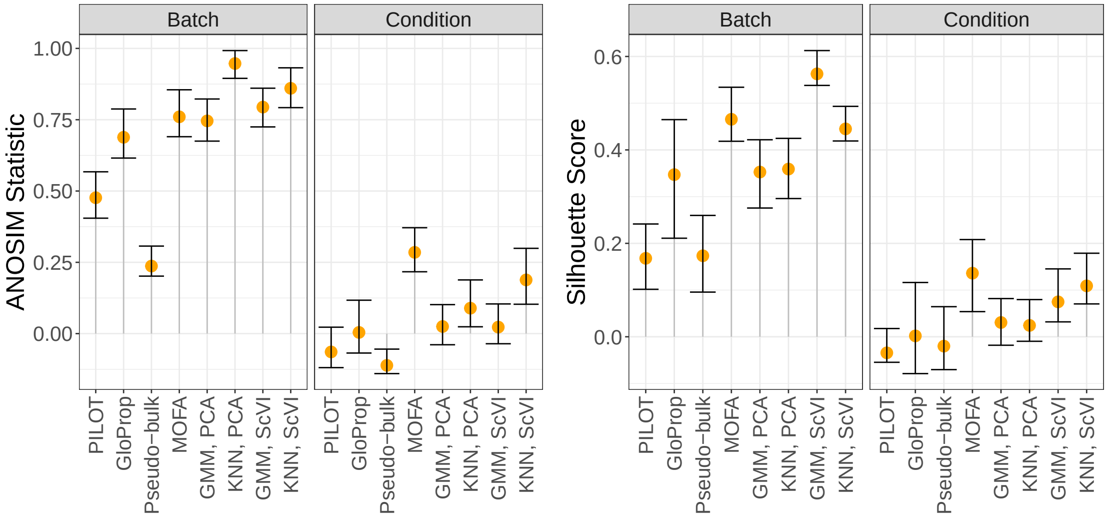


(A) ANOSIM

(B) Silhouette

Fig. S29: **Separation of different sample-level methods on COVID PBMC study [5].** The separation of samples in different batches or biological conditions based on the (A) ANOSIM Statistic and (B) Average Silhouette Width. The orange point is the value of the statistic calcualted by the indicated method, along with bootstrap confidence intervals.

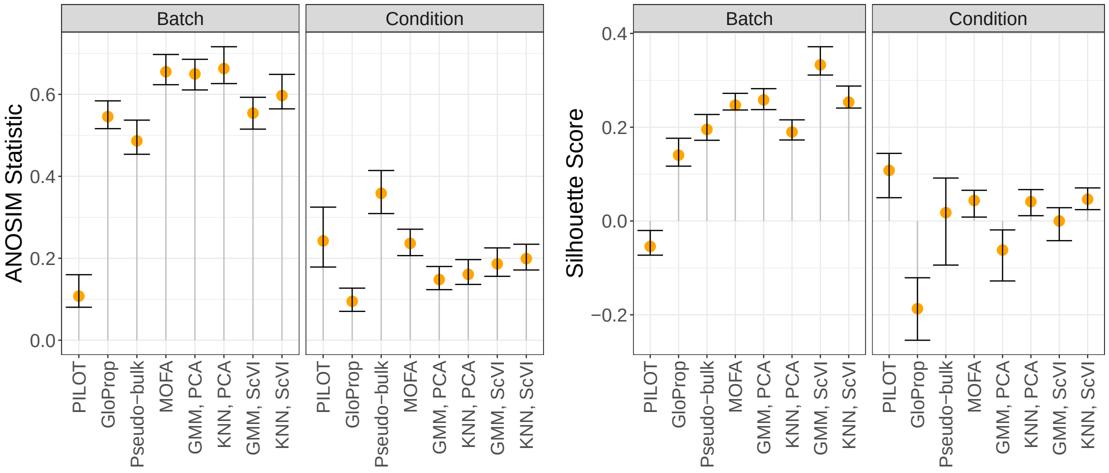


(A) ANOSIM

(B) Silhouette

Fig. S30: **Separation of different sample-level methods on on Lupus PBMC study [7].** The separation of samples in different batches or biological conditions based on the (A) ANOSIM Statistic and (B) Average Silhouette Width. Orange point is the value of the statistic calcualted by the indicated method, along with bootstrap confidence intervals.
